# Surprising Features of Nuclear Receptor Interaction Networks Revealed by Live Cell Single Molecule Imaging

**DOI:** 10.1101/2023.09.16.558083

**Authors:** Liza Dahal, Thomas GW Graham, Gina M Dailey, Alec Heckert, Robert Tjian, Xavier Darzacq

**Author notes:** Correspondence: LD, XD.

## Abstract

Type 2 Nuclear Receptors (T2NRs) require heterodimerization with a common partner, the Retinoid X Receptor (RXR), to bind cognate DNA recognition sites in chromatin. Based on previous biochemical and over-expression studies, binding of T2NRs to chromatin is proposed to be regulated by competition for a limiting pool of the core RXR subunit. However, this mechanism has not yet been tested for endogenous proteins in live cells. Using single molecule tracking (SMT) and proximity-assisted photoactivation (PAPA), we monitored interactions between endogenously tagged retinoid X receptor (RXR) and retinoic acid receptor (RAR) in live cells. Unexpectedly, we find that higher expression of RAR, but not RXR increases heterodimerization and chromatin binding in U2OS cells. This surprising finding indicates the limiting factor is not RXR but likely its cadre of obligate dimer binding partners. SMT and PAPA thus provide a direct way to probe which components are functionally limiting within a complex TF interaction network providing new insights into mechanisms of gene regulation in vivo with implications for drug development targeting nuclear receptors.

## Introduction

Complex intersecting regulatory networks govern critical transcriptional programs to drive various cellular processes in eukaryotes. These networks involve multiple transcription factors (TFs) binding shared cis-regulatory elements to elicit coordinated gene expression (Gerstein et al., 2012; Pan et al., 2009; Reményi et al., 2004). A distinct layer of combinatorial logic occurs at the level of specific TF–TF interactions, which can direct TFs to distinct genomic sites. Thus, fine tuning of TF-TF interactions can modulate TF-gene interactions to orchestrate differential gene expression. Often, dimerization between different members of a protein family can generate TF-TF combinations with distinct regulatory properties resulting in functional diversity and specificity (Nandagopal et al., 2022; Puig-Barbé et al., 2023). For example, E-box TFs of the basic helix-loop-helix (bHLH) family such as MYC/MAD share a dimerization partner MAX. Several studies have shown that switching of heterocomplexes between MYC/MAX and MAD/MAX results in differential regulation of genes and cell fate (Amati et al., 1993; Bouchard et al., 2001; Hurlin and Huang, 2006; Xu et al., 2001).

Type-2 nuclear receptors (T2NRs) present another classic example of such a dimerization network wherein the distribution of heterodimeric species regulates gene expression (Bwayi et al., 2022; Chan and Wells, 2009; Chen et al., 2018; Wang et al., 2017). T2NRs constitute an extensive group of basic leucine zipper (bZIP) TFs that share a common modular structure composed of a well-conserved ligand binding domain that mediates heterodimerization with their obligate partner Retinoid X receptor (RXR) and a highly conserved DNA binding domain (DBD) that recognizes consensus sequences termed direct response elements (DREs) (Evans and Mangelsdorf, 2014). Unlike Type 1 nuclear receptors, T2NRs do not depend on ligand binding for DNA engagement. Rather, binding of ligands to the ligand binding domain (LBD) of chromatin-associated T2NR heterodimers results in eviction of co-repressors and recruitment of co-activators (Evans and Mangelsdorf, 2014; McKenna and O’Malley, 2002) (Figure 1A).

**Figure 1:**
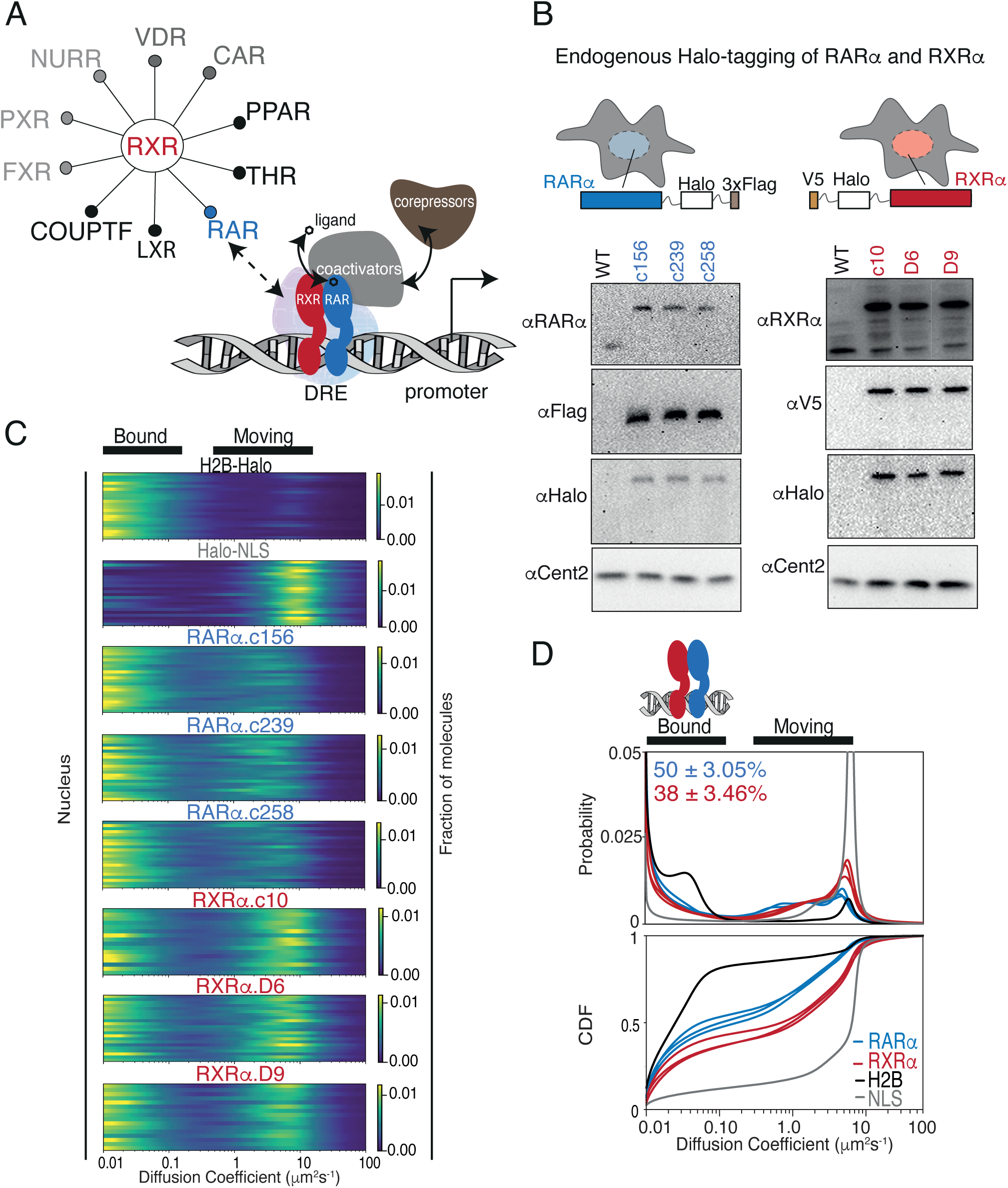
Endogenous Halo-tagging of RARα RXRα to characterize their diffusive behaviour. **(A)** Schematic showing Type II nuclear receptors (T2NRs) like RXR-RAR bind direct response elements (DREs) as heterodimers to activate or repress transcription by recruiting coactivators (in presence of ligand) or coreporessors (in absence of ligand). A competitive interaction network between the obligate heterodimeric partner RXR with other T2NRs acts as a complex regulatory node for gene expression. Mechanastic features of protein-protein interaction within this regulatory node and its affect on chromatin binding in live cells is yet to be explored. **(B)** Cartoon showing Halo-tagging scheme of RARα and RXRα alongwith western blots of U2OS wild-type (WT) and knock-in (K.I) RARα (left) and RXRα (right) homozygous clones. **(C)** Fast single molecule tracking (fSMT). Likelihood of diffusion coefficients based on model of Brownian diffusion with normally distributed localization error for H2B-Halo (black), Halo-NLS (grey), RARα clones (blue) and RXRα clones (red) with black lines on top of the figure illustrating bound and moving polulations. Each line represents a nucleus. **(D)** Diffusive spectra, probability density function (top) and cumulative distribution function (CDF) (bottom) -with drawing illustrating bound states as heterodimers of RARα and RXRα bound to chromatin.

As the common obligate partner of many other TFs, MAX and RXR are thought to act as the ‘core’ for their respective dimerization networks. It has been postulated that availability of such core TFs is likely to be limited in cells, resulting in competition between the various partner TFs involved in the network (Chan and Wells, 2009; Walker et al., 2005). Indeed, early studies using purified proteins revealed that MAD and MYC compete for binding to MAX with equal affinities, and reduced complex formation was seen for either heterodimer with increasing amounts of a competing partner (Ayer et al., 1993; Baudino and Cleveland, 2001). Similarly, the T2NRs Liver X receptor (LXR) and Peroxisome proliferator activated receptor (PPAR) were observed in vitro to have reduced binding to their respective response elements in the presence of competing T2NRs (Ide et al., 2003; Matsusue et al., 2006; Yoshikawa et al., 2003).

Whether such in vitro systems would capture the complexity and competitive dynamics at play in live cells has remained an unresolved issue. Moreover, to date, in vivo studies of competitive heterodimerization networks have not examined TFs at endogenous expression levels and thus may not accurately recapitulate their dynamic interactions with each other and with chromatin (Fadel et al., 2020; Grinberg et al., 2004) or do not account for expression levels of relevant TFs in individual cells due to the population averaging nature of most such studies (Rehó et al., 2023) (Reho 2023). To overcome these potential shortcomings, we employed fast single-molecule tracking (fSMT) (Boka et al., 2021; Dahal et al., 2023; Elf et al., 2007; Hansen et al., 2018, 2017) and its newly developed complement, proximity-assisted photoactivation (PAPA-SMT) (Graham et al., 2022) to test the effects of varying the stoichiometry of a core TF (RXRα) and its partner TF (RARα). We find that, contrary to expectations, the core component in cancer cells is not limiting but rather is in sufficient excess to accommodate more RARα even in the presence of many other endogenous partner T2NRs.

## Results

### Live-cell SMT of knock-in Halo-tagged RAR and RXR

As an initial test case for studying T2NR interactions, we focused on the heterodimeric partners RARα and RXRα, which are endogenously expressed alongside various other T2NRs in U2OS cells, a well-established cancer cell line for single-molecule tracking (Hansen et al., 2017; McSwiggen et al., 2019) (Figure 1A, Table S1). Using CRISPR/Cas9-mediated genome editing, we generated clonal lines with homozygous knock-in (KI) of HaloTag at the N-terminus of RXRα and the C-terminus of RARα (Heckert et al., 2022; Los et al., 2008) (Figure S1A). Western blotting confirmed that RARα and RXRα were tagged appropriately and expressed at similar levels to the untagged proteins (Figure 1B), while co-IP experiments verified that Halo-tagged RARα and RXRα heterodimerize normally as expected (Figure S1B). In addition, we confirmed using luciferase assays that the RAR ligand, all-trans retinoic acid (atRA), activated retinoic acid responsive element (RARE)-driven gene expression in wild-type and homozygously edited clones, confirming the normal transactivation function of the tagged proteins (Figure S1C). Confocal live cell imaging of cells stained with Janelia Fluor X 549 (JFX549) Halo ligand displayed normal nuclear localization of both Halo-tagged RARα and RXRα (Figure S1D).

To evaluate how RARα and RXRα explore the nuclear environment and interact with chromatin, we used fSMT with a recently developed Bayesian analysis method, SASPT (Heckert et al., 2022), to infer the underlying distribution of diffusion coefficients within the molecular population, yielding a “diffusion spectrum” (Figure 1C and D). From these diffusion spectra we can extract quantitative parameters of subpopulations (peaks) such as mean diffusion coefficients and fractional occupancy, allowing us to measure the chromatin-bound fraction (*f*_bound_) in live cells (Figure 1C and D). To benchmark our SMT measurements, we first compared the diffusion spectra of RARα-Halo and Halo-RXRα to that of H2B-Halo (which as expected is largely chromatin-bound) and Halo-3×NLS (which is mostly unbound). We observed that RARα and RXRα both exhibit a clearly separated slow diffusing population (< 0.15 μm^2^s^−1^) along with a faster mobile population (1-10 μm^2^s^−1^) (Figure 1C). The former we classify as ‘bound’ since it represents molecules diffusing at a rate indistinguishable from that of Halo-H2B (chromatin motion). The proportion of molecules diffusing at <0.15 μm^2^s^−1^ is henceforth designated as *f*_bound_. In comparison with Halo H2B (*f*_bound_ = 75.5 ± 0.7%) and Halo-NLS (*f*_bound_ = 10 ± 0.9%), RARα and RXRα have intermediate levels of chromatin binding (*f*_bound_ of 50 ± 3.0% and 38 ± 3.5%, respectively) (Figure 1D). The *f*_bound_ was reproducible between three clonal cell lines of RARα (49± 1%, 47± 1%, 53± 1%) and RXRα (36 ± 1%, 36±1%, 42±1%). Finally, although recent studies have reported an increase in chromatin interaction upon agonist treatment (Brazda et al., 2014, 2011; Rehó et al., 2023, 2020), we did not observe a significant change in *f*_bound_ of RARα and RXRα upon atRA treatment, nor did atRA treatment appear to alter the fast-moving populations of RARα and RXRα (Figure S3A & B). These results are consistent with the classic model in which dimerization and chromatin binding of T2NRs are ligand independent.

### Chromatin binding of RARα and RXRα can be saturated in live cells

Chromatin binding by an individual TF within a dimerization network is predicted to be sensitive to its expression level (Klumpe et al., 2023). To test how the expression levels of RARα and RXRα affect *f*_bound_, we overexpressed Halo fusions from stably integrated transgenes in U2OS cells (Figure 2A). After confirming transgene expression by western blotting (Figure 2A), we compared the abundance of endogenous and exogenous Halo-tagged RARα and RXRα using flow cytometry (Cattoglio et al., 2019) (Figure S4A). Average cellular abundance of endogenous RARα and RXRα obtained from biological replicates of each homozygous clone were similar (Figure 2B). In contrast, expression of exogenous RARα-Halo was approximately four times that of the endogenous protein, while expression of Halo-RXRα was nearly twenty times that of endogenous RXRα (Figure 2B). Chromatin binding of overexpressed RARα was reduced by approximately half compared to endogenous (*f*_bound_ = 27±0.72%) (Figure 2C & S4B), while that of overexpressed RXRα was decreased even more dramatically to 11±0.75%—a value barely above the *f*_bound_ of the Halo-NLS control (Figure 2C & S4B). Using nuclear fluorescence intensity as a rough proxy for protein concentration in individual cells, we observed a negative correlation in single cells between TF concentration and *f*_bound_ (Figure S4C). These results imply that chromatin binding of both RARα and RXRα in U2OS cells is saturable—that is, the total number of chromatin-bound molecules does not increase indefinitely with expression level but is in some way limited.

**Figure 2:**
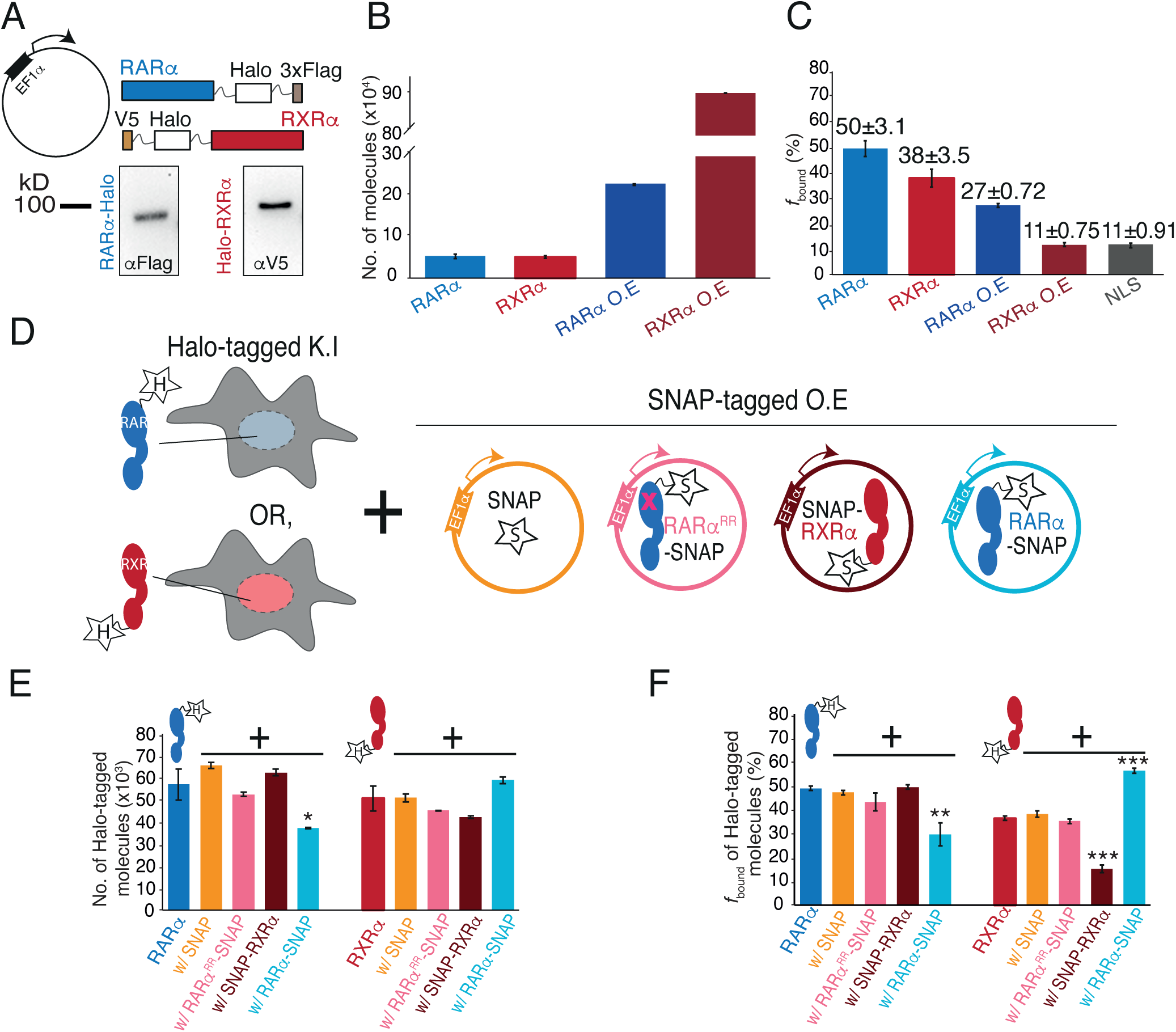
Chromatin binding of RARα and RXRα can be saturated and is limited by RARα. **(A)** Schematic and western blot of stably integrated EF1α promoter driven Halo-tagged (HT) RARα (left) and RXRα (right) overexpression in WT U2OS cells. **(B)** Bar plot; y-axis shows number of Halo-tagged (HT) knock-in (K.I) and overexpressed (O.E) RARα (blue) and RXRα (red) molecules quantified using flow cytometry. **(C)** Bar plot; y-axis depicts chromatin bound fraction (*f*_bound_%) of K.I and O.E RARα, RXRα compared to Halo-NLS (control). **(D)** Assay condition schematics to determine which of the partners in the RARα/RXRα hetero-dimer complex is limiting for chromatin binding; parental K.I HT RARα or RXRα clones with O.E SNAP (orange), SNAP-RXRα (brown), RARα-SNAP (light blue), and RAR^RR^α-SNAP (pink) using stably integrated EF1α promoter driven transgene. **(E)** Bar chart; y-axis denotes number of K.I HT RARα and RXRα molecules (depicted as blue and red cartoon respectively with ‘H’ labelled star attached) in presence or absence of trans-gene products. Error bars denote stdev of the mean from three biological replicates. **(F)** Bar plot showing *f*_bound_% of K.I HT RARa and RXRa in presence or absence of exogenously expresssed SNAP proteins. Error bars for (B), (E) denote stdev of the mean from three biological replicates. Error bars for (C), (F) represent stdev of bootstr-apping mean. P value ≤ 0.001(***), ≤ 0.01(**) & ≤ 0.05 (*).

### RARα limits chromatin binding of RXRα

We next assessed how overexpression of RARα and RXRα affects *f*_bound_ of the endogenous proteins by stable integration of SNAP-tagged RARα or RXRα transgenes in Halo-KI RARα and RXRα cell lines (Figure 2D). As controls, we also stably integrated transgenes expressing SNAP-NLS or a SNAP-tagged dimerization-incompetent RARα (RARα^RR^) (Bourguet et al., 2000; Zhu et al., 1999) (Figure 2D). We validated disruption of the RARα^RR^-RXRα interaction using Rosetta modelling (Shringari et al., 2020) and co-immunoprecipitation (co-IP) (Figure S5A). Using flow cytometry, fluorescent gels, and western blots we first assessed if transgene expression alters expression of endogenous Halo-tagged RARα and RXRα (Figure 2E and S5B-D). While no drastic changes in the cellular abundance of KI Halo-tagged RARα or RXRα was observed in the presence of SNAP, SNAP-RXRα or RARα^RR^-SNAP proteins, the abundance of KI Halo-tagged RARα was approximately halved when RARα-SNAP was overexpressed (Figure 2E and S5B). This is likely distinct from previously reported ligand dependent RARα degradation (Tsai et al., 2023; Zhu et al., 1999) (see Discussion).

We then carried out a series of fSMT experiments to understand how *f*_bound_ of the endogenous RARα or RXRα is altered when its binding partner is present in excess (Figure S7A and S7B). Surprisingly, we found that the *f*_bound_ of endogenous RARα-Halo (47±1%) was largely unchanged upon expression of RXRα-SNAP (49±1%), consistent with the control SNAP (47±1%) (Figure 2F and S7B), implying that RARα *f*_bound_ is not limited by the availability of RXR. In contrast, RARα-SNAP expression significantly decreased chromatin binding of endogenous RARα-Halo to 29±5% (Figure 2F and S7B). However, overexpression of mutant RARα^RR^-SNAP did not change the *f*_bound_ of endogenous RARα (43±4%) (Figure 2F and S7B), suggesting that heterodimerization with RXRα is required for this effect.

In the reciprocal experiment with endogenous Halo-RXRα, the initial *f*_bound_ (35±1%) was barely altered by overexpression of SNAP (38±1%) or RARα^RR^-SNAP (35±1%). As expected, overexpression of SNAP-RXRα reduced the *f*_bound_ of endogenous Halo-RXRα to 16±2%. In contrast to RARα, however, overexpression of RARα-SNAP increased the *f*_bound_ of endogenous Halo-RXRα to 56±1%. This result suggests that endogenous RXRα chromatin binding is limited by the availability of RAR (Figure 2F and S7B), and not due to limiting amounts of the universal dimer core partner RXR as would expected based on current models in the literature.

While it may seem paradoxical that RAR is limiting for RXR binding, given the similar number of molecules of endogenous RARα and RXRα per cell (Figure 2B), this is likely due to the presence of other endogenous RXR paralogs (See Table S1 and Discussion). It is also probable that some fraction of RXR binding to chromatin arises from other T2NRs that can produce chromatin binding competent dimers with RXR (Figure 1A, Table S1 and Discussion). Notwithstanding these complexities, the results of the above SMT measurements in the presence of varying amounts of partner TFs allows us to infer that endogenous RXRα is likely in excess while RXR partners are limiting in U2OS cells.

### PAPA-SMT shows that RARα-RXRα dimerization correlates with chromatin binding

The above results support a model in which the total RXR pool (including all paralogs) is in stoichiometric excess of its partners in U2OS cells. To more directly monitor RXR-RAR interactions, we employed a recently developed SMT assay by our lab, proximity-assisted photoactivation (PAPA) (Graham et al., 2022). PAPA detects protein-protein interactions by using excitation of a “sender” fluorophore with green light to reactivate a nearby “receiver” fluorophore from a dark state (Graham et al., 2022) (Figure 3A). As an internal control, violet light is used to induce direct reactivation (DR) of receiver fluorophores independent of their proximity to the sender (Figure 3A). See our supplementary note for a more detailed description of the steps involved in a PAPA-SMT experiment.

**Figure 3.**
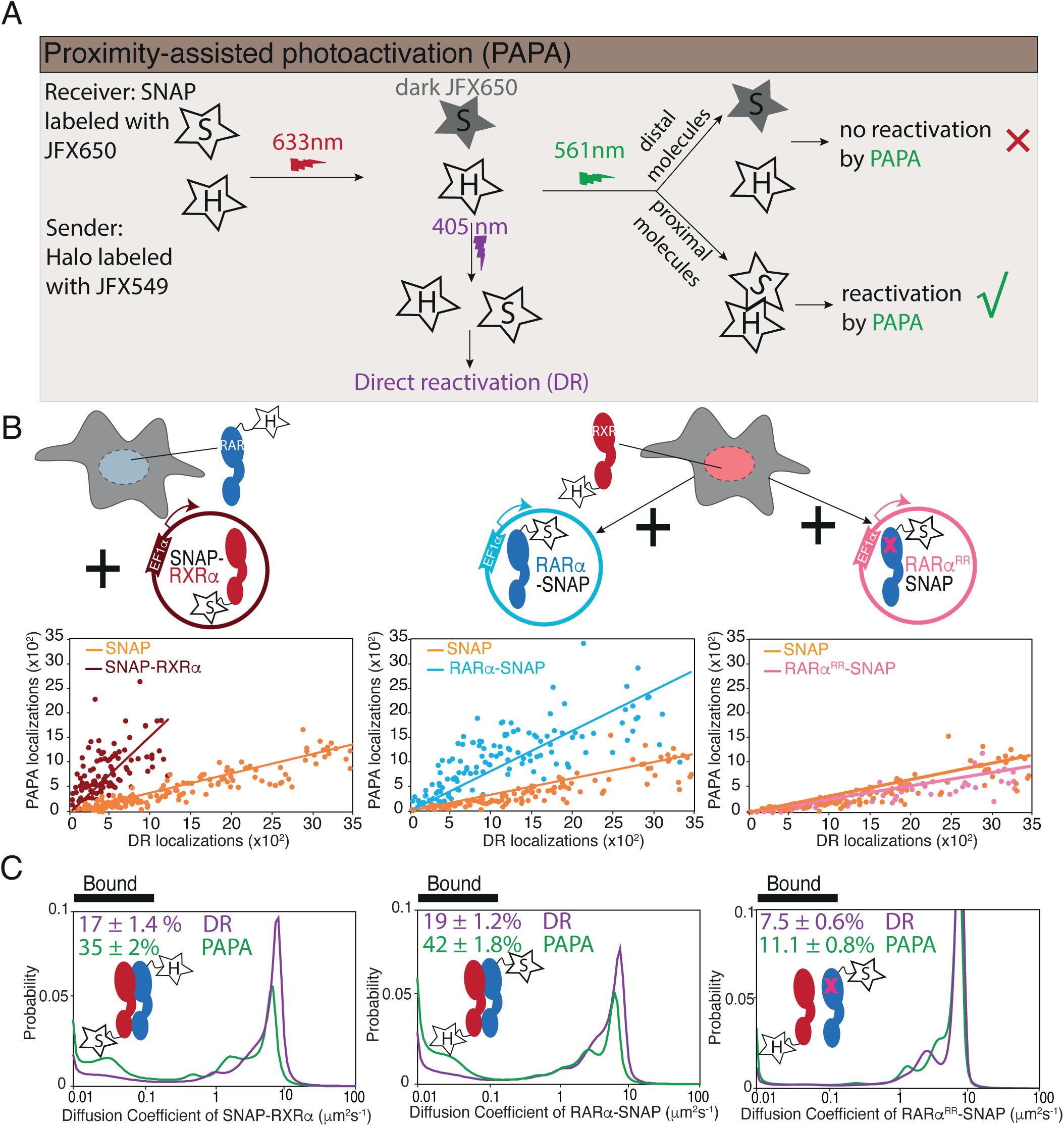
PAPA-SMT shows direct interaction between Halo-tagged K.I and SNAP-tagged overexpressed RARα and RXRα in live cells. **(A)** Schematic illustrates how PAPA signal is achieved. Firstly, SNAP-tagged(ST) protein is labelled with ‘receiver’ fluorophore like JFX650 (star with letter ‘S’) and Halo-tagged (HT) protein is labelled with ‘sender’ fluorophore like JFX549 (star with letter ‘H’). When activated by intense red light the receiver fluorophore goes into a dark state (grey star with S). Upon illumination by green light, the receiver and sender molecules distal to one another do not get photoactivated (red X) but receiver SNAP molecules proximal to the sender gets photoactivated (green *√*). Pulses of violet light can induce direct reactivation (DR) of receiver independent of interaction with the sender. PAPA experiments, **(B)** Plots showing PAPA versus DR reactivation, **(C)** Diffusion spectra of PAPA and DR trajectories obtained for ST proteins for the represented conditions; parental HT RARα knock-in (K.I) cell, stably expressing RXRα-SNAP (brown, left panel), as well as parental HT RXRα K.I cell stably expressing RARα-SNAP (light blue, middle panel) and RARα^RR^-SNAP (light pink, right panel). A linear increase in PAPA versus DR reactivation is seen for non-interacting SNAP controls and a sublinear increase is seen for interacting SNAP proteins. Respective colored lines show linear fits of the data (see residuals in Figure S8B). SNAP control data are replotted in middle and right panels. Cartoon inside diffusion spectra depicts if the expressed HT and ST proteins are expected to interact or not. *f*_bound_ errors represent stdev of bootstrapping mean.

We applied PAPA-SMT to cell lines expressing SNAP-tagged RARα and RXRα transgenes in the background of Halo-tagged endogenous RXRα or RARα, respectively (Figure 3B). As controls, we also imaged SNAP-3×NLS and RARα^RR^-SNAP, which are not expected to interact with Halo-RARα or RXRα. First, we plotted the number of single-molecule localizations reactivated by green and violet light in each cell (Figure 3B). As expected, a low level of background reactivation by green light was observed for SNAP negative control, reflecting the baseline probability that an unbound SNAP molecule will at some low level be close enough to Halo to observe PAPA (Figure 3B, orange points) (Graham et al., 2022). Violet light induced DR and green light induced PAPA were linearly correlated as expected, since both are proportional to the number of receiver molecules. However, a greater PAPA signal was seen for the Halo-RXRα → RARα-SNAP and RARα-Halo → SNAP-RXRα combinations than for the negative control, consistent with direct protein-protein interactions (Figure 3B). The relation between DR and PAPA was sub-linear (Figure 3B, left and middle panel, see residuals of the linear fit in Figure S8B), consistent with saturation of binding to the Halo-tagged component. In contrast, the ratio of PAPA to DR for Halo-RXRα → RARα^RR^-SNAP was similar to the SNAP negative control, confirming that PAPA signal depends on a functional RAR-RXR interaction interface (Figure 3B, right panel).

Next, we compared the diffusion spectra of molecules reactivated by DR and PAPA (Figure 3C). For both Halo-RXRα → RARα-SNAP and RARα-Halo → SNAP-RXRα combinations, PAPA trajectories had a substantially higher chromatin-bound fraction than DR trajectories, indicating that a greater proportion of SNAP-tagged RARα/RXRα binds chromatin when it is in complex with its Halo-tagged partner. Only a slight shift in bound fraction was seen for RARα^RR^-SNAP (*f*_bound,DR_ = 7.5±0.6%; *f*_bound,PAPA_ = 11.1±0.8%), comparable to that seen for the SNAP negative control (*f*_bound,DR_ = 7.1±0.5% and *f*_bound,PAPA_ = 11.2±0.9% for Halo-RXRα; *f*_bound,DR_ = 7.3±0.4% and *f*_bound,PAPA_ = 9.3±0.7 for RARα-Halo; (see Discussion), confirming that RARα^RR^ fails to form chromatin binding-competent heterodimers with RXR.

The enrichment of chromatin-bound molecules in the PAPA-reactivated population of both RAR and RXR is consistent with heterodimerization mediating chromatin binding, that is further validated by the low *f*_bound_ of the dimerization-incompetent mutant.

## Discussion

According to the current consensus models for how nuclear receptor interaction networks operate, specific T2NRs members compete for a limited pool of the “core” partner RXR in cells, thus establishing a competitive regulatory network (Chan and Wells, 2009; Fadel et al., 2020; Rehó et al., 2023; Wang et al., 2005; Wood, 2008; Yoshikawa et al., 2003). However, previous in vivo studies were carried out with overexpressed proteins, limiting their ability to accurately address this model of competition (Fadel 2019, Reho 2023). Here we used SMT in live cells with endogenously tagged proteins and carefully controlled protein levels of two players, RARα and RXRα in the T2NRs dimerization network, thereby directly addressing a fundamental question - whether RAR (partner) or RXR (core) is limiting for chromatin association.

In stark contrast to the generally accepted T2NR competition model, our results reveal that in U2OS cells, formation and chromatin binding of RAR-RXR heterodimers are limited by the concentration of RAR and not the core subunit RXR (Figure 4). PAPA-SMT directly confirmed in vivo that the association with RXRα promotes chromatin binding of RARα and vice versa (Figure 3C). However, unexpectedly, overexpression of RARα increases *f*_bound_ of endogenous RXRα, while the reverse is not true (Figure 2F), indicating that RXR is not limiting but is rather constrained by the availability of its binding partners. Note that even though the number of RXRα and RARα molecules is about the same and the heterodimer complex has a predicted 1:1 stoichiometry (Rastinejad, 2022; Rastinejad et al., 2000), the *f*_bound_ of endogenous RARα (50%) and RXRα (38%) differed by about 10% (Figure 1D and 2B). This surprising result could be explained by expression of additional RXR isoforms (most likely RXRβ) in U2OS cells (Figure 1A, Table S1, Figure S2B). The fact that the *f*_bound_ of overexpressed RARα decreases only two-fold, even though it is in four-fold excess over endogenous RXRα, likewise suggests that there are other heterodimerization partners of RARα available (Figure 2A and 2B, see Table S1). Moreover, the *f*_bound_ of overexpressed RXR is essentially the same as the NLS control, indicating this excess core partner remains nearly totally unbound (Figure 2C). This is consistent with endogenous RXR already being in excess such that any additional RXR would remain mostly monomeric and unbound to chromatin. Hence, despite not directly measuring the protein levels of all RXR isoforms, we can still deduce in U2OS cells that RXR is in excess relative to its binding partners.

**Figure 4.**
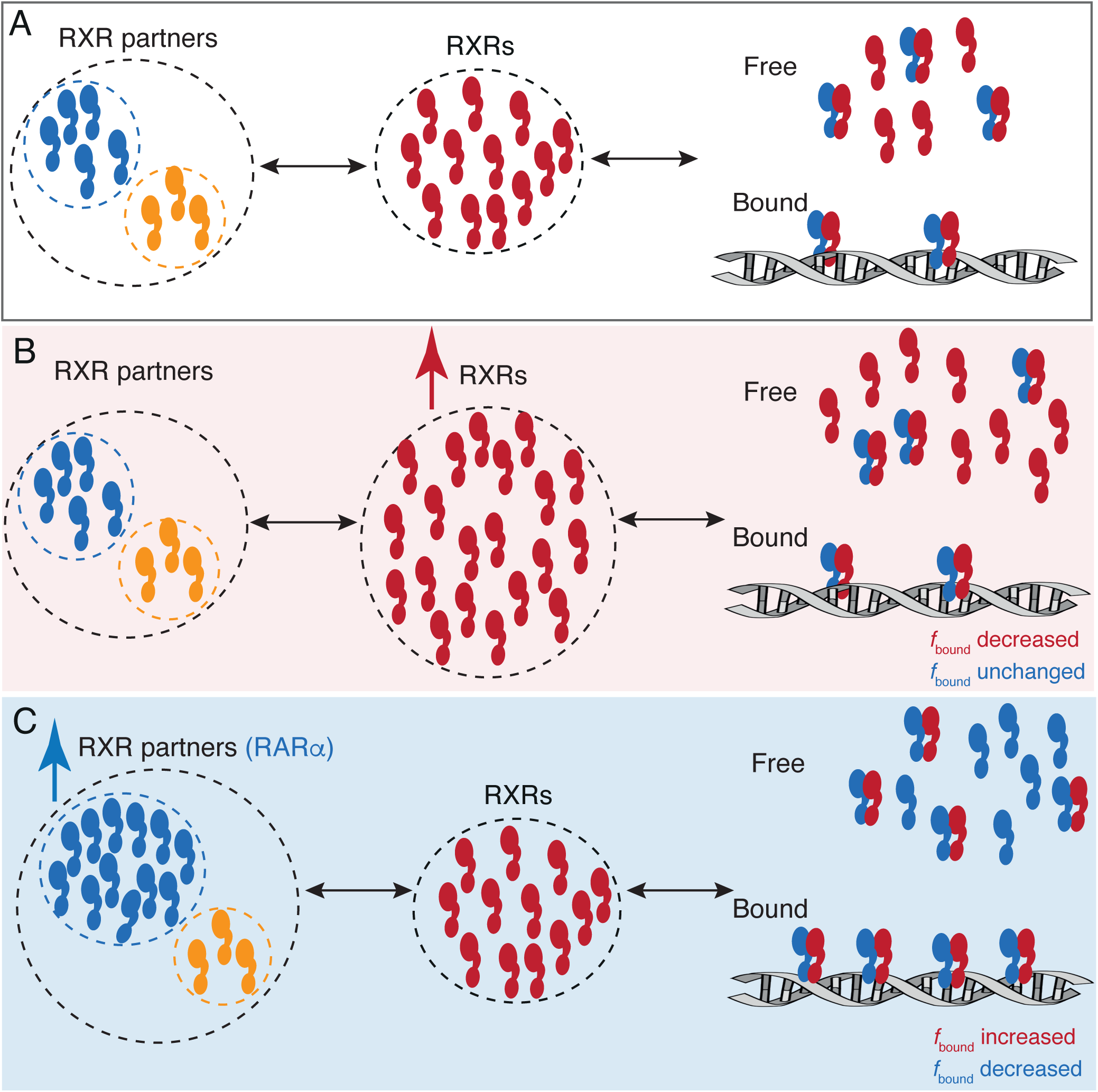
A model for RARα limited chromatin binding of RARα-RXRα heterodimers. **(A)** Pool of RXRα (red) and RXR partners (RARα – blue, other T2NRs-yellow) along with some number of chromatin bound RARα-RXRα heterodimers exist under normal conditions. **(B)** When the pool of free RXRα is increased, the number of chromatin bound RARα-RXRα heterodimers does not change. **(C)** When the pool of RARα is increased, chromatin binding RARα-RXRα heterodimers increases, until it reaches saturation. Note: For simplicity we have omitted to show heterodimerization of other T2NRs (yellow) with RXRα (red).

Control of RAR-RXR heterodimer concentration by RAR abundance makes sense considering the observation that RAR protein levels appear to be regulated by multiple feedback mechanisms: First, we find that overexpression of RARα significantly lowers the expression of endogenous RARα (Figure 2E) implying either that RAR participates in negative autoregulation at the transcriptional level or, that an excess of RAR causes instability at the protein level. We favor the latter possibility because overexpression of RARα reduces not only the expression of endogenous RARα but also its *f*_bound_ (Figure 2F), indicating a reduction in dimerization and chromatin binding. It is possible that endogenous RAR may be more readily degraded when not bound by RXR, analogous to what was recently shown for the c-MYC/MAX heterodimers (Mark et al., 2023). Second, as has been previously reported, RARα expression decreases upon ligand treatment (Ismail and Nawaz, 2005; Kopf et al., 2000; Osburn et al., 2001; Tsai et al., 2023; Zhu et al., 1999). Curiously, a 50% reduction in RARα upon addition of ligand did not affect the *f*_bound_ of either RARα or RXRα (Figure S3A and S3B), suggesting that most chromatin binding of endogenous RXRα in U2OS cells depends on heterodimerization partners other that RARα (see Table S1).

In contrast to the generally accepted model, we thus envision a network of T2NR heterodimers not always driven by competition for the core TF partner (RXR). Instead, an excess of RXR may ensure independent regulation of the different T2NRs without disrupting crosstalk between them (Figure 4).

In this first study, we have not examined other T2NRs expressed in U2OS cells or whether there is a similar excess of RXR in other cell types. Therefore, we cannot rule out that competition for RXR between T2NRs occurs in some cases since different cell types express RXR and partner T2NRs in varying amounts. It seems likely that chromatin binding by any given T2NR will depend on the concentrations of all other T2NRs. It is thus important to determine how interactions within the dimerization network are perturbed upon up or down-regulation of one or more T2NR, especially since dysregulation of T2NR expression has been reported in several types of cancers as well as other diseases (Brabender et al., 2005; Collins-Racie et al., 2009; Frigo et al., 2021; Long and Campbell, 2015). Here we have shown that SMT and PAPA-SMT provide one empirical way to determine which set of components in a dimerization network is stoichiometrically limiting in a given cell type, without having to measure the concentration of every T2NR or the affinity of every interaction. The basic framework we have established here could also be extended to probe the effect of ligands or small molecules on the T2NR interaction network or other dimerization networks seen in bHLH or leucine zipper family of transcription factors, providing useful information about critical regulatory network interactions often implicated in diseases.

## Methods

### Cell culture and stable cell line generation

U2OS cells were grown in Dulbecco’s modified Eagle’s medium (DMEM) with 4.5 g/L glucose supplemented with 10% fetal bovine serum (FBS) (HyClone, Logan UT, Cat. #AE28209315), 1 mM sodium pyruvate (Thermo Fisher 11360070), L-glutamine (Sigma #G3126-100G), Glutamax (ThermoFisher #35050061) and 100 U/ml penicillin-streptomycin (Thermo Fisher #15140122) at 37°C and 5% CO_2_. Cells were subcultured at a ratio of 1:4 to 1:10 every 2 to 4 days for no longer than 30 passages. Regular mycoplasma testing was performed using polymerase chain reaction (PCR). Phenol containing media was used for regular cell culture and Phenol red-free DMEM (Thermo Fisher #21063029) supplemented with 10% FBS and 100 U/ml penicillin-streptomycin was used for imaging.

Stable cell lines expressing the exogenous gene products (Table S2) were generated by PiggyBac transposition and antibiotic selection. Gibson assembly was used to clone genes of interest into a PiggyBac vector containing a puromycin or neomycin resistant gene. Plasmids were purified by Zymo midiprep kit (Zymo D4200) and all cloning was confirmed by Sanger sequencing. Cells were transfected by nucleofection using the Lonza Cell line Nucleofector Kit V (Lonza, Basel, Switzerland, #VVCA1003) and the Amaxa Nucleofector II device. For each transfection cells were plated 1-2 days before nucleofection in a 6 well plate until they reach 70-90% confluency. Cells were trypsinized, resuspended in DMEM media and centrifuged at 200xg for 2 mins before the media was aspirated. Cells were then resuspended in 100 ul Lonza transfection reagent (82 μl Kit V solution + 16 μl of Supplement #VVCA1003) containing 0.4 μg of SuperPiggyBac transposon vector and 0.8 μg of donor PiggyBac plasmid and transferred to an electroporation cuvette. Cells were electroporated using program X-001 on the Amaxa Nucleofector II (Lonza). Transfected cells were cultured in DMEM growth media without any antibiotics for 24-48 hrs and then selected for 10 days with 1 μg/ml puromycin (Thermo Fisher #A1113803) or 1mg/ml neomycin (G418 Sulfate, Thermo Fisher #10131027). After selection polyclonal cell lines were maintained in the selection media containing required antibiotics.

For ligand treatment, 100μM all trans retinoic acid (atRA) stock was prepared by dissolving atRA powder (CAS No: 302794, Sigma Aldrich #R2625) in Dimethyl sulfoxide (DMSO) (Sigma Aldrich #D2650) and was diluted 1:100,000 or 1:1000 in growth media to final concentration of 1 nM or 100 nM respectively. The same volume of DMSO used for 100 nM atRA treatment (0.1%) is used for the control group as a condition without atRA treatment. Cells were treated 24 hr in either atRA or DMSO alone before imaging.

### Genome editing cell lines

Knock-in (K.I) cell lines were generated as previously described (Hansen 2017) with some modifications. Halo-tagging of endogenous RARα was described in our previously published work (Heckert et al., 2022) which we have further validated for the current study. For Halo-tagging we designed sgRNAs using CRISPOR web tool (Concordet and Haeussler, 2018). Since exon 1 of RXRα is very short (only 9 aa) we chose to gene edit at the start of exon 2 for a successful tagging. sgRNAs were cloned into the Cas9 plasmid (a gift from Frank Xie) under the U6 promoter (Zhang Lab) with an mVenus reporter gene under the PGK promoter. Repair vectors were cloned in a basic pUC57 backbone for N-terminal tagging and pBluescript II SK (+) (pBSKII+) backbone for C-terminal tagging, with 500 bp left and right homology arms on either side of the Halo-tag sequence. Two guide/repair for N-terminal and three guide/repair vector pairs for C-terminal were attempted; only-N-terminal clones were ultimately recovered. Each sgRNA/donor pair were transfected to approximately 1 million early passage U2OS cells, at 1:3 ratio of sgRNA/donor (Total 5 μg DNA) and plated in a 6 well plate. 48 hrs after transfection, Venus-positive cells were FACS sorted and cultured for another 7-10 days. Then, Halo-positive cells (stained with TMR) were sorted individually into single wells of 96 well plates and cultured for another 12-14 days. Clones were expanded and genotyped using PCR. PCR was done using one primer upstream of the left homologous arm and the other primer downstream of the right homologous arm. Another PCR with either external primer paired with a corresponding internal primer located in the Halo-Tag coding region was done for further validation. Homozygous clones with the correct genotype (V5-Halo-RXRα clones C10, D6 and D9) were confirmed by Sanger sequencing and western blotting.

### Antibodies

The following antibodies were used for western blotting: mouse monoclonal anti-RARα [H1920] (Abcam, #ab41934) diluted at 1:400, rabbit monoclonal anti-RXRα [EPR7106] (Abcam, #ab125001) diluted at 1:500, rabbit polyclonal anti-RXRα (Proteintech, #212181AP), mouse monoclonal anti-V5 tag (Thermo Fisher, #R960-25) diluted at 1:5000, mouse monoclonal anti-FLAG [M2] (Sigma Aldrich, #F1804) diluted at 1:5000, mouse monoclonal anti-Halo (Promega, #G9211) diluted at 1:500, mouse monoclonal anti-TBP (Abcam, #ab51841) diluted at 1:5000, rabbit polyclonal anti-Centrin 2 (Proteintech, # 158771AP) diluted at 1:1000, goat polyclonal anti-mouse IgG light chain specific (Jackson ImmunoResearch, # 115035174) diluted at 1:10000, mouse monoclonal anti-rabbit IgG light chain specific (Jackson ImmunoResearch, #211032171) diluted at 1:10000.

The following antibodies were used for co-immunoprecipitation: rabbit polyclonal anti-V5 (Abcam, # ab9116) for immunoprecipitation, mouse monoclonal anti-FLAG [M2] (Sigma Aldrich, #F1804) diluted at 1:5000 and mouse monoclonal anti-V5 tag (Thermo Fisher, #R960-25) diluted at 1:5000 for blotting.

### Western blotting

For western blots, cells growing in 6 well or 10 cm plates at 80-90% confluency were scraped and pelleted in ice cold phosphate-buffered saline (PBS) containing protease inhibitors. Cell pellets were resuspended using 500-1000 μl hypotonic buffer (100 mM NaCl, 25 mM HEPES, 1 mM MgCl_2_, 0.2 mM EDTA, 0.5% NP-40 alternative) containing protease inhibitors-1× aprotinin (Sigma, #A6279, diluted 1:1000), 1 mM benzamidine (Sigma, #B6506), 0.25 mM PMSF (Sigma #11359061001) and 1× cOmplete EDTA-free Protease Inhibitor Cocktail (Sigma, #5056489001) alongwith 125 U/ml of benzonase (Novagen, EMD Millipore #71205-3). Resuspended cells were gently rocked at 4°C for atleast 2 hrs after which 5 M NaCl was added. Cells were left rocking for extra 30 mins at 4°C and then centrifuged at max speed at 4°C. Supernatants were quantified using Bradford and 15-20 μg was loaded on 8% Bis-Tris SDS gel. Wet transfer to 0.45 μm nitrocellulose membrane (Thermofisher #45004031) was performed in a transfer buffer (15 mM Tris-HCl, 20 mM glycine, 20% methanol) for 60-80 mins at 100 V, 4°C. Membranes were blocked in 10% non-fat milk in 0.1% TBS-Tween (TBS-T) for 1 hr at room temperature (RT) with agitation. Membranes were then blotted overnight with shaking at 4°C with primary antibodies diluted in TBS-T with 5% non-fat milk. After 5× 5 mins washes in 0.1% TBS-T, membranes were incubated at RT with HRP conjugated light chain secondary antibodies diluted 1:10000 in TBS-T with 5% non-fat milk, for 1 hr with agitation. After 5× 5 mins washes in 0.1% TBS-T, membranes were incubated for 2 mins in freshly prepared Perkin Elmer LLC Western Lightning Plus-ECL, enhanced Chemiluminescence Substrate (Thermo Fisher, #509049326). Finally, membranes were imaged with BioRad Chemidoc imaging system (BioRad, Model No: Universal Hood III).

### Luciferase assays

pGL3-RARE luciferase, a reporter containing firefly luciferase driven by SV40 promoter with three retinoic acid response elements (RAREs), a gift from T. Michael Underhill (Addgene plasmid 13456; http://n2t.net/addgene:13458; RRID:<u>Addgene_13458</u>; <u>Hoffman et al., 2006</u> was used for luciferase assays. A pRL-SV40 vector (Promega #E2261) expressing Renilla luciferase with SV40 promoter was used as a control to normalize luciferase activity. Cells were plated on 6 well plate at least 48 hrs before being co-transfected with 200 ng pGL3-RARE-luciferase and 10ng Renilla luciferase vector, using TransIT-2020 Transfection Reagent (Mirus Bio, #MIR5404). A day after transfection, cells were treated with 100 nM atRA or 0.1% DMSO (control). Luciferase assay was performed the next day, using Dual-Luciferase Reporter Assay System (Promega, #E1910) according to manufacturer’s protocol, on the Glomax Luminometer (Promega). The relative luciferase activity was calculated by normalizing firefly luciferase activity to the Renilla luciferase activity to control for transfection efficiency.

### Co-Immunoprecipitation

For co-immunoprecipitation (CoIP) experiments, Cos7 cells were plated to 2× 15 cm dishes for each condition, at 60-70% confluency. DNA Lipofectamine 3000 (Thermo Fisher #L3000015) was used to co-transfect the cells with plasmids expressing RARα-Halo-3×FLAG and RXRα-V5 or V5-Halo-RXRα and RARα-3×FLAG or RAR^RR^α-Halo-3×FLAG and RXRα-V5. According to manufacturer’s instructions, 500 μl Opti-MEM medium (Thermo Fisher #31985062) was combined with 20 μl P3000 reagent and the plasmids for each condition. To control for the ratio of transfected DNA to the reagents, total transfected DNA mass was kept at 10 μg for each condition using empty pBSK vector (Addgene # 212205). 500 μl Opti-MEM medium containing 20 μl Lipofectamine 3000 reagent was subsequently added to the plasmid containing mixture and incubated at RT for 15 mins, after brief pipetting to mix the solutions. The mixture was then divided equally to the cells plated in 2× 15 cm dishes. COS7 cells were cultured in DMEM media with 4.5 g/L glucose supplemented with 10% fetal bovine serum (FBS), 1 mM sodium pyruvate, L-glutamine, Glutamax and 100 U/ml penicillin-streptomycin and maintained at 37°C and 5% CO_2_.

Cells were collected from plates 48 hrs after transfection by scraping in ice-cold 1× PBS with protease inhibitors (1× aprotinin, 0.25 mM PMSF, 1 mM benzamidine and 1× cOmplete EDTA-free Protease Inhibitor Cocktail). Collected cells were pelleted, flash frozen in liquid nitrogen and stored at −80°C. On the day of CoIP experiments, cell pellets were thawed on ice, resuspended to 700 μl of cell lysis buffer (10 mM HEPES pH 7.9, 10 mM KCl, 3 mM MgCl_2_, 340 mM sucrose (1.16gr) and 10% glycerol) with freshly added 10% Triton X-100. Cells were rocked at 4°C for 8 mins to allow lysis and centrifuged for 3 mins at 3000 ×*g*. The cytoplasmic fraction was removed, and the nuclear pellets were resuspended in 1 ml hypotonic buffer (100 mM NaCl, 25 mM HEPES, 1 mM MgCl_2_, 0.2 mM EDTA, 0.5% NP-40 alternative) containing protease inhibitors and 1 μl benzonase. After rocking for 2-3 hrs at 4°C, the salt concentration was adjusted to 0.2 M NaCl final and the lysates were rocked for another 30 mins at 4°C. Supernatant was removed after centrifugation at maximum speed at 4°C for 20 mins and quantified by Bradford. Typically, 1 mg of protein was diluted in 1 ml 0.2 mM CoIP buffer (200 mM NaCl, 25 mM HEPES, 1 mM MgCl_2_, 0.2 mM EDTA, 0.5% NP-40 alternative) with protease inhibitors and cleared for 2 hrs at 4°C with magnetic Protein G Dynabeads (Thermo Fisher, #10009D) before overnight immunoprecipitation with 1 μg per 250 μg of proteins of either normal serum IgGs or specific antibodies as listed above. Some precleared lysates were kept overnight at 4°C as input. Magnetic Protein G Dynabeads were also precleared overnight with 0.5% BSA at 4°C. Next day the precleared Protein G dynabeads were added to the antibody containing samples and incubated at 4°C for 2 hrs. Samples were briefly spun down and placed in a magnetic rack at 4°C for 5 mins to remove the CoIP supernatant. After extensive washes with the CoIP buffer, the proteins were eluted from the beads by boiling for 10 mins in 1× SDS-loading buffer and analyzed by SDS-PAGE and western blot.

### Flow cytometry

Cells were grown in 6 well dishes. On the day of the experiment, cells were labeled with 500 nM Halo-TMR for 30 mins, followed by one quick wash with 1× PBS and 15 mins wash in dye free DMEM before trypsinization. Cells were pelleted after centrifugation, resuspended in fresh medium, filtered through 40 μm filtration unit, and placed on ice until fluorescence read out by Flow Cytometry (within 30 mins). Using a LSR Fortessa (BD Biosciences) flow cytometer, live cells were gated using forward and side scattering and TMR fluorescence emission read out was filtered using 610/20 band pass filter after excitation with 561 nm laser. Mean fluorescence intensity of the samples and absolute abundance were calculated as described in Cattoglio et al 2019, using a previously quantified Halo-CTCF cell line as a standard (Cattoglio et al., 2019).

### Cell preparation and dye labeling for imaging

For confocal imaging ∼50,000 cells were plated in tissue culture treated 96 well microplate (Perkin Elmer, #6055300) a day before imaging. Cells were labeled with Halo and/SNAP for 1 hr and incubated in dye-free media for 20 mins after a quick 1× PBS wash to remove any free dye. Phenol free media containing Hoechst (1 μM) was added to the cells and incubated for 1 hr before proceeding with imaging.

For SMT, 25 mm circular No. 1.5H precision cover glass (Marienfield, Germany, 0117650) were sonicated in ethanol for 10 mins, plasma cleaned then stored in 100% isopropanol until use. ∼250,000 U2OS cells were plated on sonicated, and plasma cleaned coverglass placed in 6 well plates. For ligand treatment, cells were treated with DMSO (control, 0.1%), 1 nM atRA or 100 nM atRA, a day after plating cells in the coverslip and, a day before imaging. After ∼24-48 hrs, cells were incubated with 100 nM PA-JFX549 dye in regular culture media for 30 mins, followed by 4× 30 mins incubations with dye-free culture media at 37°C. A quick 2× PBS wash was interspersed between each 30 mins dye-free media incubation. After the final wash, coverslips with plated cells facing upwards, were transferred to Attofluor Cell Chambers (Thermo Fisher, # A7816), and phenol free media added for imaging. For Halo-tagged RARα and RXRα homozygous clones that were stably integrated with SNAP-tagged RARα, RXRα or control transgenes under EF1α promoter; cells were double-labeled with 100 nM HaloTag ligand (HTL) PA-JFX549 and SNAP tag ligand (STL) 50 nM SF-650 simultaneously. The 30 mins labeling step was followed by the same wash steps to remove free dye as described above, before transferring the coverslips to Attofluor Cell Chambers and imaging.

For PAPA-SMT experiments, polyclonal U2OS cells stably expressing SNAP-tagged RARα or RXRα under EF1α promoter within the endogenous Halo-tagged RARα and RXRα homozygous clones were used. ∼ 400,000 polyclonal cells were plated in glass-bottom dishes (MatTEK P35G-1.5-20-C) and stained overnight with 50 nM JFX549 HTL and 5 nM JFX650 STL in phenol red-free DMEM (Thermo Fisher #21063029). Next day, cells were briefly rinsed twice with 1× PBS, incubated twice for 30 mins in phenol red-free DMEM to remove free dye, and exchanged into fresh phenol red-free medium more before imaging.

### Confocal imaging

For confocal imaging, endogenously Halo-tagged cells were labeled with Halo ligand JFX549 (100 nM) and Hoechst (1.6 μM); endogenously Halo-tagged cells overexpressing SNAP-tagged proteins were labeled with Halo ligand JFX549 (100 nM), SNAP ligand SF650 (25 nM) and Hoechst (1.6 μM). Imaging was performed at the UC Berkeley High Throughput Screening facility on a Perkin Elmer Opera Phenix equipped with 37°C and 5% CO_2_, using a built-in 40× water immersion objective.

### Live cell Single molecule tracking (SMT)

All SMT experiments were performed using a custom-built microscope as previously described (Hansen et al., 2017). Briefly, a Nikon TI microscope was equipped with a 100×/NA 1.49 oil immersion total internal reflection fluorescence (TIRF) objective (Nikon apochromat CFI Apo TIRF 100× Oil), a motorized mirror, a perfect Focus system, an EM-CCD camera (Andor iXon Ultra 897), laser launch with 405 nm (140 mW, OBIS, Coherent), 488 nm, 561 nm and 639 nm (1W, Genesis Coherent) laser lines, an incubation chamber maintaining a humidified atmosphere with 5% CO_2_ at 37°C. All microscope, camera and hardware components were controlled through the NIS-Elements software (Nikon).

For tracking endogenous as well as overexpressed Halo-tagged proteins, PA-JFX549 labelled cells were excited with 561 nm laser at 2.3 kW/cm^2^ with single band-pass emission filter (Semrock 593/40 nm for PA-JFX549). All imaging was performed using highly inclined optical sheet (HILO) illimitation (Tokunaga et al., 2008). Low laser power (2-3%) was used to locate and focus cell nuclei. Region of interest (ROI) of random size but with maximum possible area was selected to fit into the interior of the nuclei. Before tracking, 100 frames of continuous illumination with 549 laser (laser power 5%) at 80 ms per frame was recorded to later calculate mean intensity of each nuclei. After partial pre-bleaching (only for overexpressed Halo-tagged proteins), movies were taken with 1 ms pulses of full power of 561 nm illumination at the beginning of frame interval with camera exposure of 7 ms/frame, while the 405 nm laser (67 W/cm^2^) was pulsed during ∼447 μs camera transition time. Total of 30,000 frames were collected. 405 nm intensity was manually tuned to maintain low density (1 to 5 molecules fluorescent particles per frame).

For tracking experiments with endogenous Halo-tagged proteins in presence of transgenic SNAP protein products, cells were dual labelled with both PA-JFX549 and STL-SF650. Before tracking cells were excited with a 639 nm laser at 88 W/cm^2^ and single band-pass emission filter set to Semrock 676/37 nm. Low laser power (2-3%) was used to locate and focus cell nuclei. As before, ROI of random size but with maximum possible area was selected to fit into the interior of the nuclei. The emission filter was switched to Semrock 593/40 nm band-pass filter for PA-JFX549, keeping the TIRF angle, stage XYZ position and ROI the same. Cells were then excited with 549 nm and single molecules were tracked for 30,000 frames as described above.

Note, we estimated that to calculate *f*_bound_ of RARα and RXRα with minimal bias and variation we need to analyze at least 10,000 pooled trajectories from n ≥ 20 cells (Figure S1E). Therefore, all analysis of fSMT to calculate *f*_bound_ or *f*_bound_ % presented in this paper are done accordingly. At least 8-10 movies were collected for each sample as one technical replicate on a given day. Two technical replicates on two separate days were collected to produce the reported results. As observed previously for SMT experiments (McSwiggen et al., 2024) the variance within collected data was largely due to cell-to-cell variance (Figure S1E).

### PAPA-SMT

PAPA-SMT imaging was performed using the same custom-built microscope described above using an automated system (Graham et al., 2024; Walther et al., 2024). Briefly, the microscope described above was programmed using Python and NIS Elements macro language code to raster in a grid over the coverslip surface, acquire an image in the JFX650 channel, identify cell nuclei, reposition the stage to center it on a target nucleus, resize the imaging region of interest to fit the nucleus, pre-bleach JFX650 with red light (639 nm) for 5 seconds, and finally perform a PAPA-SMT illumination sequence. The illumination sequence consisted of 5 cycles of the following, at a frame rate of 7.48 ms/frame:

1. Bleaching: 200 frames of red light (639 nm), not recorded
2. Imaging: 30 frames of red light, one 2-ms stroboscopic pulse per frame, recorded
3. PAPA: 5 frames of green light (561 nm), not recorded
4. Imaging: 30 frames of red light, one 2 ms stroboscopic pulse per frame, recorded
5. Bleaching: 200 frames of red light, not recorded
6. Imaging: 30 frames of red light, one 2 ms stroboscopic pulse per frame, recorded
7. DR: 1 frames of violet light (405 nm), one 1 ms stroboscopic pulse, not recorded
8. Imaging: 30 frames of red light, one 2 ms stroboscopic pulse per frame, recorded

Laser power densities were approximately 67 W/cm^2^ for 405 nm, 190 W/cm^2^ for 561 nm, and 2.3 kW/cm^2^ for 639 nm. This illumination sequence includes non-stroboscopic illumination periods to bleach fluorophores or place them back in the dark state (steps 1 and 5), and it records only the frames immediately before and after the PAPA and DR pulses. This is done to decrease file sizes and speed up subsequent analysis compared to our previous protocol (Graham et al., 2022).

### SMT and SMT-PAPA data processing and analysis

All SMT movies were processed using the open-source software package quot (https://github.com/alecheckert/quot) (Heckert, 2022). For SMT dataset quot package was run on each collected SMT movie using the following settings: [filter] start = 0; method = ‘identity’; chunk_size = 100; [detect] method = ‘llr’; k=1.0; w=15, t=18; [localize] method = ‘ls_int_gaussian’, window size = 9; sigma = 1.2; ridge = 0.0001; max_iter = 20; damp = 0.3; camera_gain = 109.0; camera_bg = 470.0; [track] method = ‘euclidean’; pixel_size_μm = 0.160; frame interval = 0.00748; search radius = 1; max_blinks = 0; min_IO = 0; scale – 7.0. The first 500 or 1000 frames of each movie were removed due to high localization density. To confirm that all movies were sufficiently sparse to avoid misconnections, a maximum number of 5 localizations per frame was maintained (although most frames had 1 or less). To infer the distribution of diffusion coefficients from experimentally observed trajectories, we used the state array method, which is publicly available at https://github.com/alecheckert/spagl (Heckert et al., 2022). The following settings were used: likelihood_type= ‘rbme’; pixel_size_μm = 0.16; frame_interval = 0.00747; focal depth = 0.7; start_frame = 500 or 1000 frames. RBME likelihood for individual cells and occupations as the mean of the posterior distribution over state occupations marginalized on diffusion coefficient is reported. For SMT movies in Figure S4C, mean intensity of each cell was calculated using the mean gray value tool in FIJI. Mean gray value of each frame (total 100 frames) of the pre-bleaching 80ms movie was measured. Average mean gray value over the 100 frames is reported as mean intensity for each cell.

For PAPA-SMT quot package was run on each collected SMT movie using the following settings: [filter] start = 0; method = ‘identity’; chunk_size = 100; [detect] method = ‘llr’; k=1.2; w=15, t=18; [localize] method = ‘ls_int_gaussian’, window size = 9; sigma = 1.2; ridge = 0.0001; max_iter =10; damp = 0.3; camera_gain = 109.0; camera_bg = 470.0; [track] method = ‘conservative’; pixel_size_μm = 0.160; frame interval = 0.00748; search radius = 1; max_blinks = 0; min_IO = 0; scale – 7.0. Trajectories were sorted based on whether they occurred before or after green (561 nm; PAPA) or violet (405 nm; DR) reactivation pulses, and state array analysis (Heckert et al., 2022) was applied to each set of trajectories. For Figure 3C, reactivation by PAPA and DR for each cell were quantified by calculating the difference in total number of localizations before and after reactivation pulses.

### Statistical Analysis

To derive a measure of error for SMT and SMT-PAPA, we performed bootstrapping analysis on all SMT datasets. For each dataset, a random sample of size n, where n is the total number of cells in the dataset was drawn 100 times. The mean and standard deviation (stdev) from this analysis is reported.

Two tailed P values were calculated based on a normal distribution (Scipy function, scipy.stats.norm.sf) with mean equal to the difference between sample means and variance equal to the sum of the variances from bootstrap sampling (for SMT and SMT-PAPA) or biological replicates (for absolute abundance calculation of molecules from flow cytometry assay). The Šidák correction was applied to correct for multiple hypothesis testing within each experiment.

## Supporting information

Supplemental Infomration

## Acknowledgements

We would like to thank all members of the Tjian-Darzacq lab for helpful discussions and suggestions over the years, in particular Claudio Cattaglio for advice on biochemsitry experiments and Vinson Fan for suggestions on SMT data analysis. We are grateful to John J. Ferrie for aiding with Rosetta modelling as well as both him and Jonathan P. Karr for critical reading of the manuscript. We also thank Luke Lavis for providing fluorescent HaloTag ligands, CRL Flow Cytometry Facility for use of their instruments, the QB3 High Throughput Screening Facility for providing access to the Opera Phenix automated confocal microscope. This work was supported by the NIH grant U54-CA231641-01659 (to XD), and the Howard Hughes Medical Institute (to RT). TG was supported by a postdoctoral fellowship from the Jane Coffin Childs for Medical Research. AH was supported by the NIH Stem Cell Biological Engineering predoctoral fellowship T32 GM098218.

## Data Availability

Source data for Figure 1-3 are accessible through https://doi.org/10.5281/zenodo.14009060.

## Author information

**Liza Dahal**

- Li Ka Shing Center for Biomedical & Health Sciences, Department of Molecular and Cell Biology, University of California, Berkeley, Berkeley, United States
- Howard Hughes Medical Institute, University of California, Berkeley, Berkeley, United States

**Contribution:** Conceptualized, designed and executed experiments, analyzed data, validation, investigation, Writing-drafted the original manuscript, review and editing.

**Competing interests:** No competing interests declared

**Thomas Graham**

- Li Ka Shing Center for Biomedical & Health Sciences, Department of Molecular and Cell Biology, University of California, Berkeley, Berkeley, United States
- Howard Hughes Medical Institute, University of California, Berkeley, Berkeley, United States

**Contribution:** Designed, executed, and analyzed SMT-PAPA experiments, Writing-review and editing.

**Competing interests:** is an inventor of pending patent application (PCT/US2021/062616) related to the use of PAPA as a molecular proximity sensor.

**Gina M Dailey**

- Li Ka Shing Center for Biomedical & Health Sciences, Department of Molecular and Cell Biology, University of California, Berkeley, Berkeley, United States

**Contribution:** Resources.

**Competing interests:** No competing interests declared

**Alec Heckert**

- Li Ka Shing Center for Biomedical & Health Sciences, Department of Molecular and Cell Biology, University of California, Berkeley, Berkeley, United States

**Contribution:** Resources.

**Competing interests:** AH is currently an employee of Eikon Therapeutics.

**Robert Tjian**

- Li Ka Shing Center for Biomedical & Health Sciences, Department of Molecular and Cell Biology, University of California, Berkeley, Berkeley, United States
- Howard Hughes Medical Institute, University of California, Berkeley, Berkeley, United States

**Contribution:** Conceptualization, Supervision, Funding acquisition, Writing-review and editing

**Competing interests:** No competing interests declared

**Xavier Darzacq**

- Li Ka Shing Center for Biomedical & Health Sciences, Department of Molecular and Cell Biology, University of California, Berkeley, Berkeley, United States

**Contribution:** Conceptualization, Supervision, Funding acquisition, Writing-review and editing

**Competing interests:** is a co-founder of Eikon Therapeutics, Inc; is an inventor on a pending patent application (PCT/US2021/062616) related to the use of PAPA as a molecular proximity sensor.

## Notes

### Competing Interest Statement

TWG is an inventor of pending patent application (PCT/US2021/062616) related to the use of PAPA as a molecular proximity sensor.XD is a co-founder of Eikon Therapeutics, Inc.

### Summary of Updates

This version of the manuscript has been updated to add some supplemental figures, revise Figure 4, clarify a couple of text in the main manuscript as well as to correct some typographical errors.

