## Supplemental Infomration for "Surprising Features of Nuclear Receptor Interaction Networks Revealed by Live Cell Single Molecule Imaging"

### Supplementary Figures

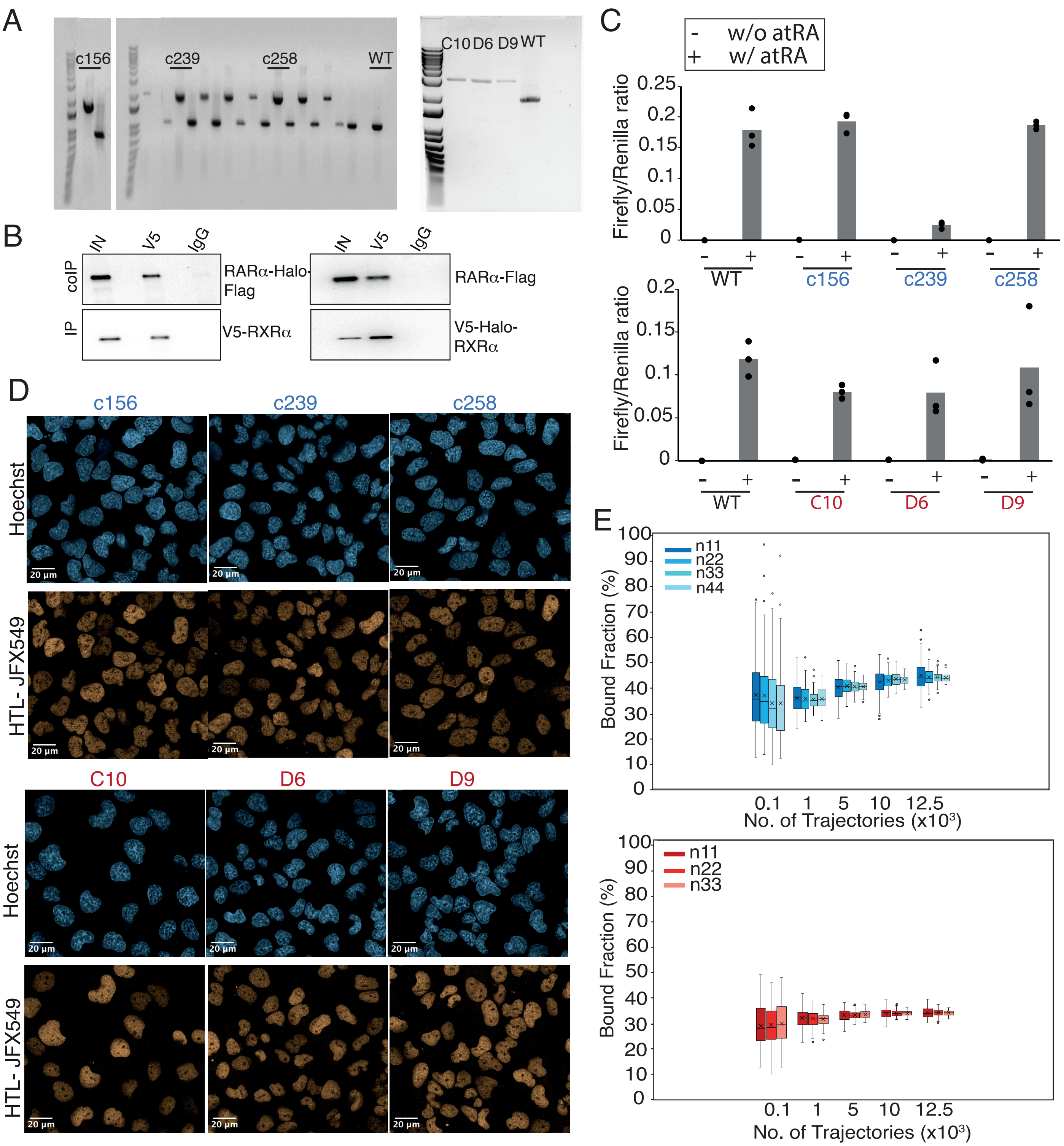


**Figure S1. (A)** Genotyping gels showing selected homozygously Halo-tagged (HT) RARα (c156, c239 & c258) and RXRα (C10, D6 & D9) clones. **(B)** Co-immunoprecipitation of over-expressed, HT RARα (left) and RXRα (right) in Cos7 cells. V5-tagged RXRα was immunoprecipitated and HT RARα was immuno-blotted (left). HT RXRα was immunoprecipitated and Flag-tagged RARα was immunoblotted (right). **(C)** Luciferase assays showing retinoic acid responsive promoter activity of wild-type (WT) and HT RARα (top) and RXRα (bottom) clones in presence and absence of all trans retinoic acid (atRA). **(D)** Confocal images of HT RARα (top) and RXRα (bottom) showing nuclear localization. **(E)** Subsampling knock-in (K.I) RARα-Halo (blue, top) and Halo-RXRα (red, bottom) trajectories from fSMT used to estimate $f_{bound}$. Sampling (with replacement) was done by number of trajectories (100, 1000, 5000, 10000 and 12500) extracted from varying number of cells (11, 22, 33 and 44). For each sample size, 100 replicates were performed and plotted as box plot showing variation and cross (x) indicating the mean.

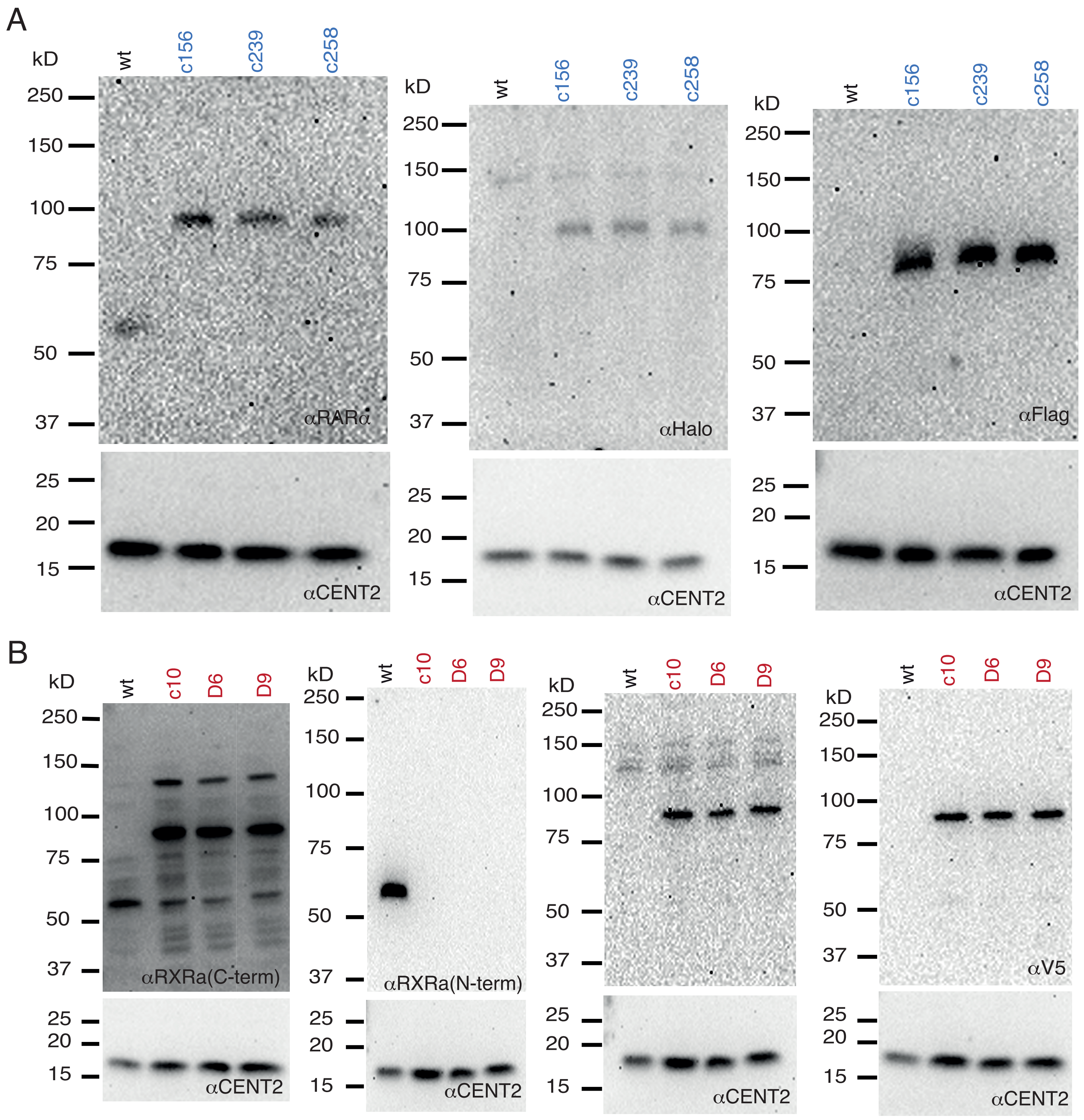


**Figure S2. (A)** and **(B)** Full length western blots of WT U2OS cells and homozygously Halo-tagged RARα and RXRα knock-in (K.I) clones. **(A)** RARα is detected in WT and K.I clones using a specific antibody against RARα. RARα is also detected in K.I clones using a specific antibody against Halo and Flag-tag. The lower part of the membrane was cut and blotted for Centrin2 as a loading control. **(B)** RXRα is detected in WT and K.I clones using a specific antibody against a C-terminal epitope of RXRα. Since this antibody also recognizes other isoforms of RXR (~54 kDa) we validated RXRα expression by using a N-term epitope specific antibody of RXRα. But this epitope sequence is disrupted by Halo-tagging at the N-terminal due to which it does not recog-nize HT-RXRα in K.I clones. Our genotyping gels (Figure S1A) give a clean single band confirming homozygous tagging and Sanger sequencing confirmed correct tagging. RXRα is also detected in K.I clones using a specific antibody against Halo and V5-tag which show a clean gel confirming correct Halo-tagged RXRα product. The lower part of the membrane was cut and blotted for Centrin2 as a loading control.

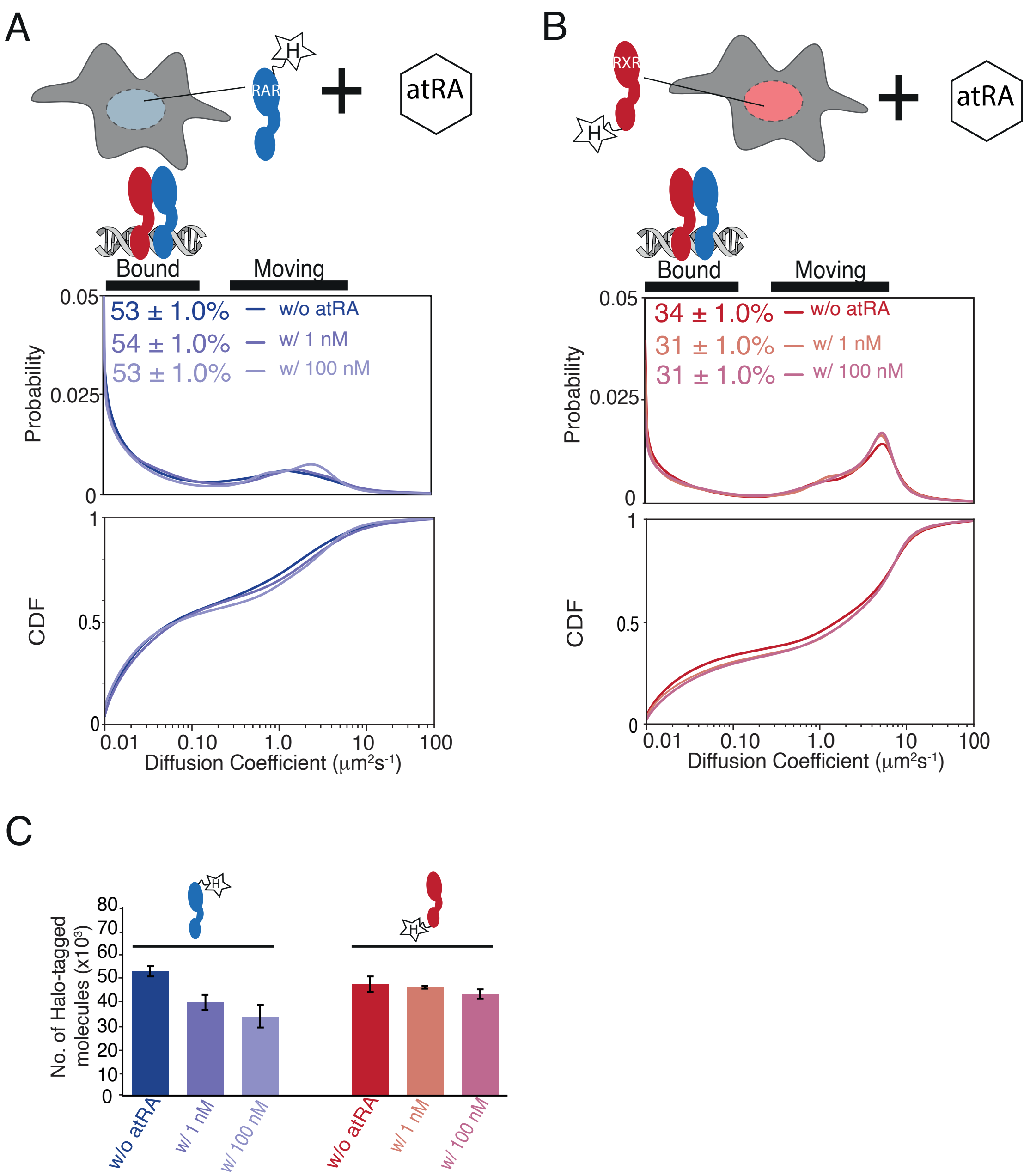


**Figure S3.** Diffusion spectra and cumulative distribution function of knock-in (K.I) (**A**) RARα-Halo and (**B**) Halo-RXRα in absence and presence of atRA. Cells were treated without or with 1 nm, and 100 nM atRA for 24 hours before SMT experiments. DMSO was added to cells without atRA treament as a control. **(C)** Cellular abundance of K.I RARα and RXRα with or without 24 hours treatment with atRA using flow cytometry analysis. Y-axis of bar plot shows number of Halo-tagged RARα and RXRα molecules. Error bars represent stdev of mean for three biological replicates.

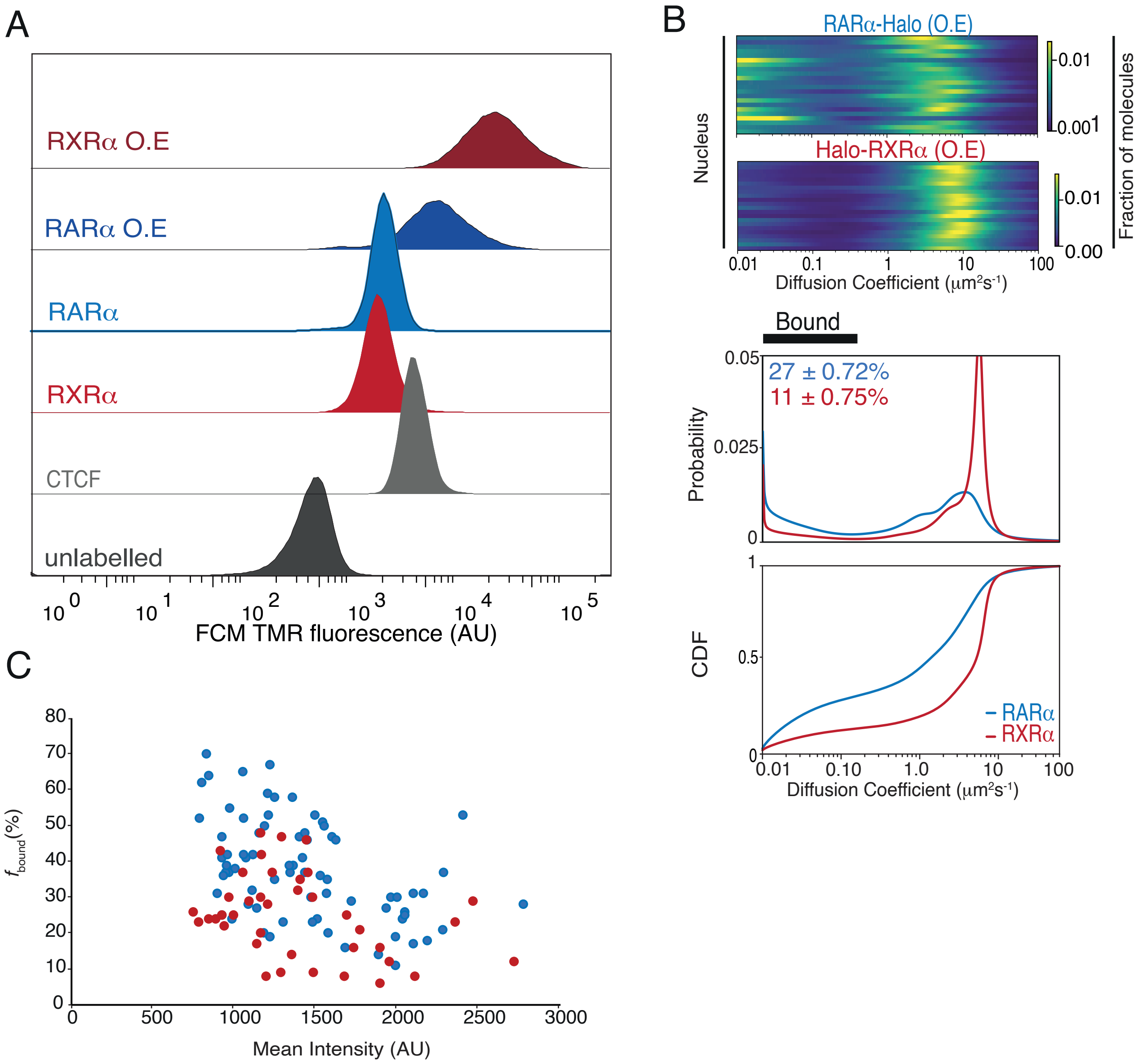


**Figure S4.** **(A)** Cellular abundance of Halo-tagged (HT) knock-in (K.I) versus overexpressed (O.E) RARα and RXRα compared with standard Halo-CTCF cell line using flow cytometry analysis. X-axis shows TMR fluorescence intensity of HT molecules in U2OS cells. **(B)** Likelihood of diffusion coefficients for O.E HT RARα and RXRα. Each line represents a nucleus. Diffusion spectra showing probability density function and cumulative distribution function for O.E HT RARα and RXRα is shown underneath. **(C)** $f_{bound}$% of K.I and overexpressed HT RARα and RXRα extracted from single cells plotted against mean nuclear intensity of single cells. Each dot represents a data collected from a single nucleus.

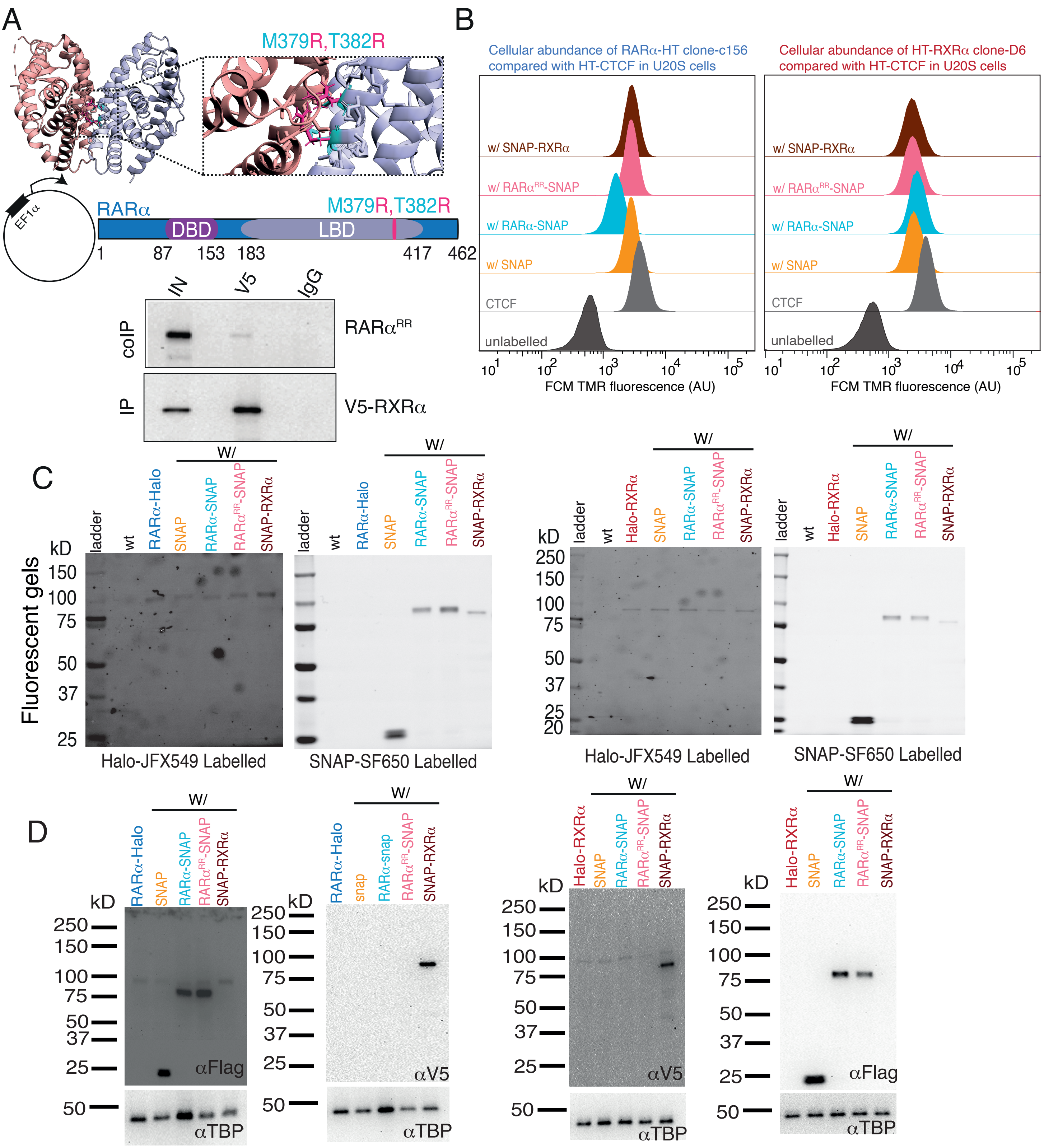

**Figure S5. (A)** Rosetta modelled structure of the ligand binding domain (LBD) of RARα (pale blue) and RXRα (pale red) heterodimer (PDB code: 1DKF) with point mutation M379R and T382R. Original residues M379 and T382 are shown as cyan sticks and substituted residue arginine (R) is show as pink sticks. Below the PDB structure is a schematic representation of M379R,T382R mutation in RARα ($RAR\alpha^{RR}$) alongwith co-immunoprecipitation in Cos7 cells. V5-tagged RXRα was immunoprecipitated and $RAR\alpha^{RR}$ was immunoblotted. $RAR\alpha^{RR}$ disrupts heterodimerization with RXRα. **(B)** Cellular abundance of knock-in (K.I) Halo-tagged (HT) RARα and RXRα in presence of overexpressed (O.E) SNAP proteins using flow cytometry (FCM) analysis. FCM estimated fluorescence of TMR labelled HT RARα and RXRα molecules were compared with Halo-CTCF in U2OS alongwith unlabelled U2OS control. **(C)** Fluorescent gels and **(D)** western blots showing HT K.I RARα and RXRα alongwith O.E SNAP proteins in stably integrated conditions. U2OS cell were labelled with Halo ligand JFX549 (100nM) and SNAP ligand SF650 (50nM) before lysis.

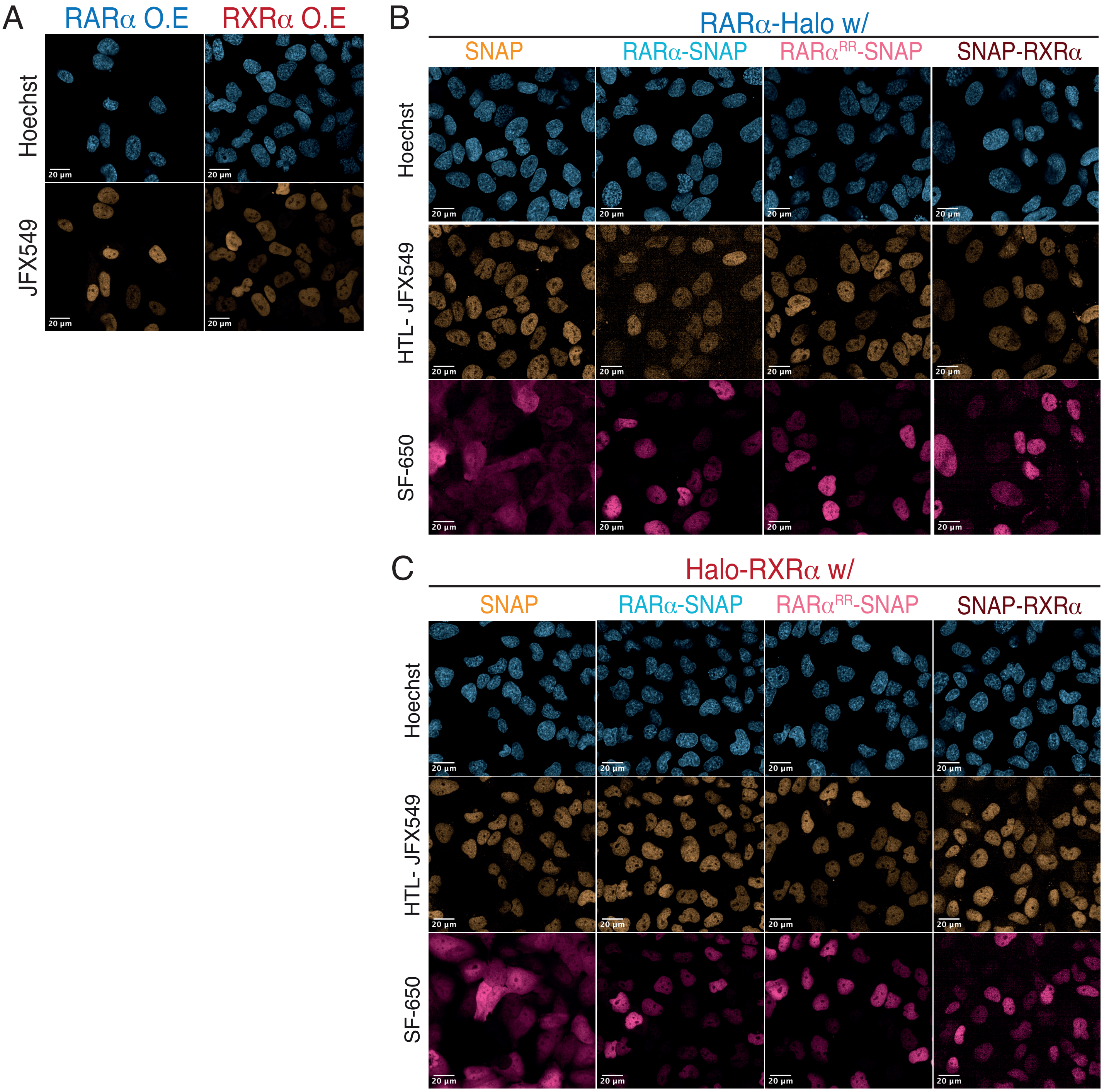


**Figure S6.** **(A)** Confocal images of overexpressed (O.E) Halo-tagged (HT) RARα and RXRα. Confocal images of U2OS cells expressing knock-in (K.I) HT **(B)** RARa and **(C)** RXRa in presence of stably integrated O.E SNAP proteins. HT molecules were labeled with 100 nM JFX549 and SNAP-tagged (ST) molecules were labeled with SF-650.

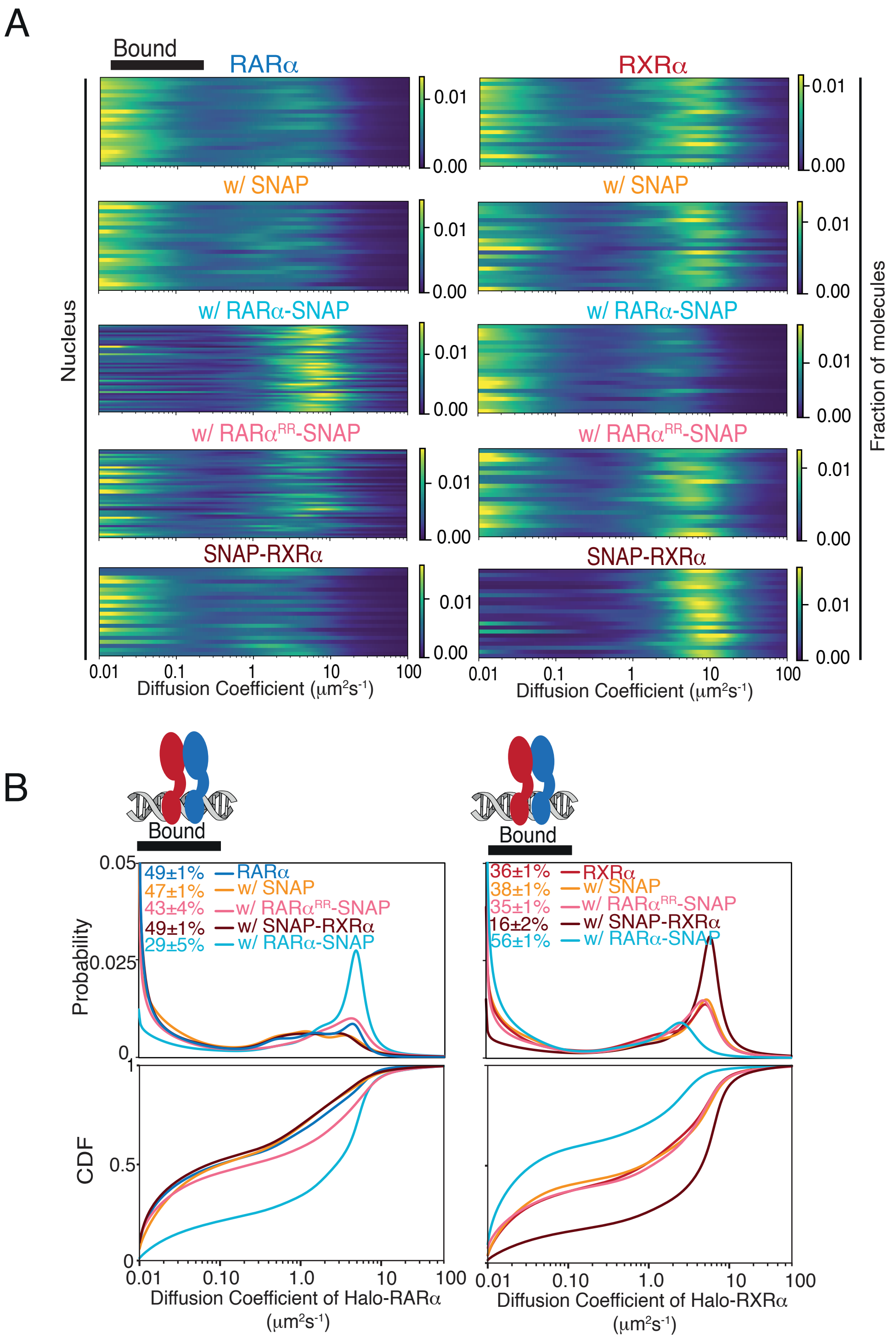


**Figure S7.** fSMT of Halo-tagged (HT) RARα and RXRα with (w/) diferent transgene products. **(A)** Likelihood diffusion coefficients and **(B)** Diffusion spectra, Probability density function (top) and cumulative distribution function (bottom) for knock-in (K.I) HT RARα (left) and RXRα (right) in absence and presence of stably integrated overexpressed (O.E) SNAP proteins. Each line in **(A)** represents a nucleus. $f_{\mathrm{bound}}$ reported shows chromatin binding of RXRα is limited by the availability of RARα but not vice versa. Cartoon above the spectra illustrates bound population as heterodimers of RARα and RXRα bound to chromatin.

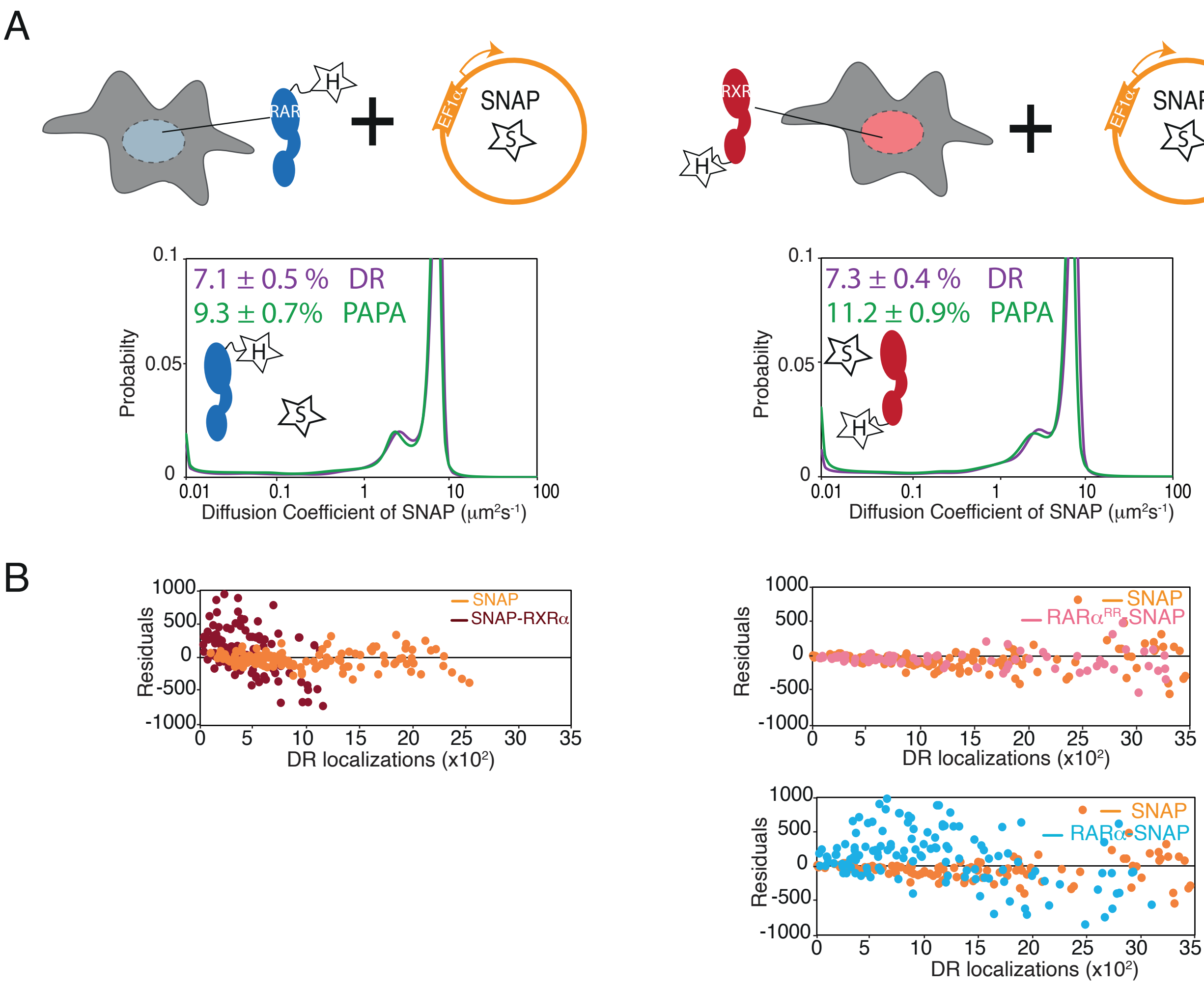


**Figure S8.** Controls for PAPA expreiments **(A)** Diffusion spectra of PAPA and DR trajectories obtained for SNAP proteins below each corresponding conditions presented as cartoons; the parental Halo-tagged (HT) RARα (left) and RXRα (right) knock-in (K.I) cells stably expressing control SNAP protein (orange). Cartoon inside diffusion spectra depicts that the expressed Halo and SNAP proteins are not expected to interact. **(B)** Residuals of the linear fit for PAPA vs DR localizations for SNAP (orange) and SNAP-RXRα (brown) proteins in presence of HT RARα (left panel), as well as SNAP (orange), $RAR\alpha^{RR}$-SNAP (light pink) and RARα-SNAP (light blue) proteins in presence of HT RXRα (right panel). Same SNAP control data are re-plotted in all plots shown in (B).

A

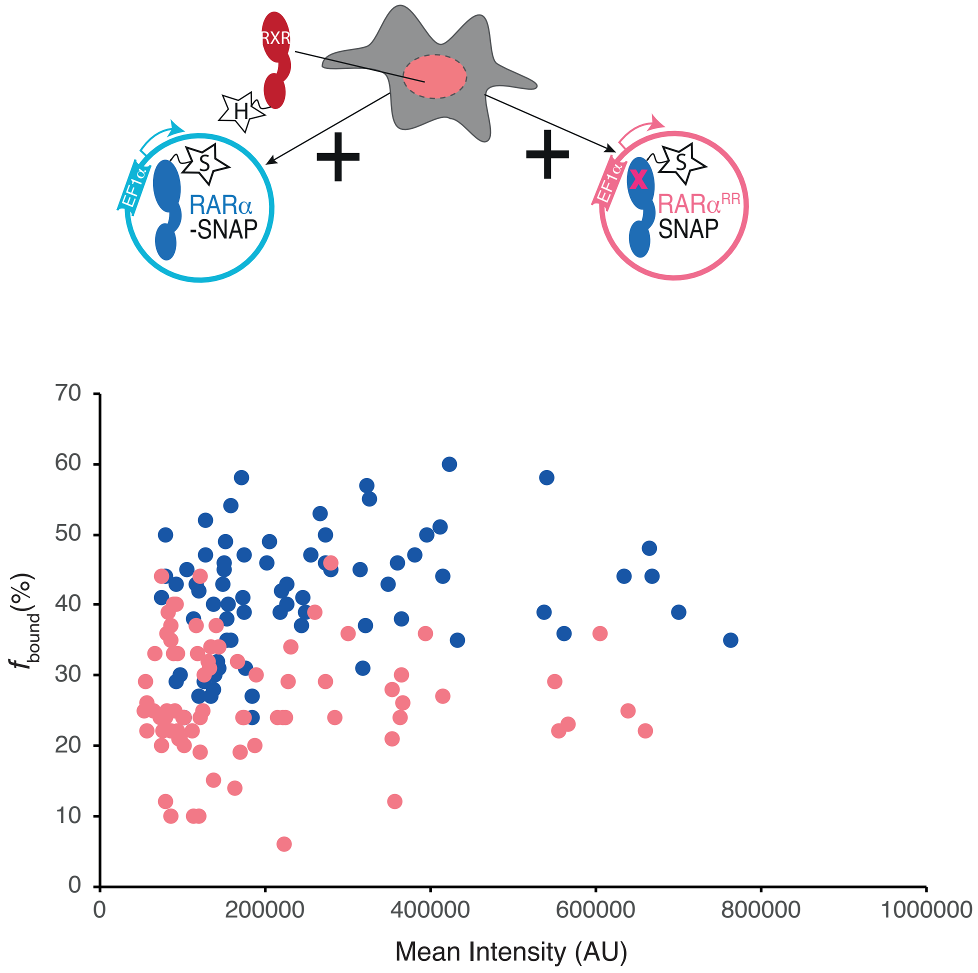


**Figure S9. (A)** $f_{\mathrm{bound}}$% of K.I HT RXRα in presence of overexpressed RARα-SNAP and RARα$^{RR}$-SNAP extracted from single cells plotted against mean nuclear intensity of single cells. Each dot represents a data collected from a single nucleus.

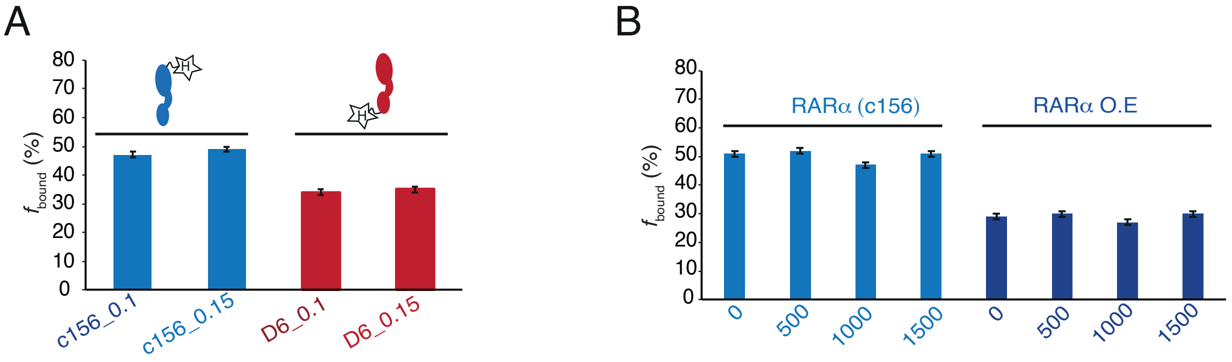


**Figure S10. (A)** Bar plot showing $f_{bound}$% of K.I HT RARα (c156) and RXRα (D6) calculated with cut-off of 0.1 (μm$^2$s$^{-1}$) and 0.15(μm$^2$s$^{-1}$) diffusion coefficients. **(B)** Bar plot showing $f_{bound}$% of K.I HT RARα (c156) and RARα O.E calculated with cut-off of 0, 500, 1000 and 1500 frames. Error bars denote stdev of bootstrapping mean.

**Supplementary Note**

Brief overview of the steps involved in a PAPA-SMT experiment:

A PAPA-SMT (Graham et al., 2022) experiment involves several steps (Fig. 3A): 1) A SNAP-tagged target protein is labeled with a "receiver" fluorophore such as JFX650, while a Halo-tagged partner protein is labeled with a "sender" fluorophore such as JFX549. 2) The JFX650 receiver is placed in a dark state by illumination with intense red (639 nm) light. 3) A pulse of green (561 nm) light is used to excite the JFX549 sender, reactivating nearby dark-state JFX650 molecules. Because association of the two proteins brings the sender and receiver together, this step selectively reactivates complexes of the two proteins. 4) After switching off the green light, red light is used to image reactivated receiver molecules. 5) As an internal control, a violet (405 nm) light pulse is used to reactivate the receiver by "direct reactivation" (DR), independent of proximity with the sender (Heilemann et al., 2008). 6) After switching off the violet light, reactivated molecules are again imaged with red light. Steps 3-6 are repeated for multiple rounds, and SMT trajectories occurring after green and violet pulses are analyzed separately using SASPT. Trajectories occurring after a green pulse (PAPA trajectories) are enriched for complexes between the SNAP- and Halo-tagged proteins, while trajectories occurring after a violet pulse (DR trajectories) approximate a random sample from the whole population.

### Supplementary Tables

**Table S1.** T2NRs expressed in U2OS cells according to the Expression Atlas (Papatheodorou et al., 2020; Prakash et al., 2023) and the Human Protein Atlas Databank (Thul et al., 2017; Uhlén et al., 2015).

| Name | TPM (Expression Atlas) | TPM (Human Protein Atlas) |
|---|---|---|
| RARα ( NR1B2) | 43 | 47.1 |
| RARβ (NR1B2) | 0.6 | 0.1 |
| RARγ (NR1B3) | 39 | 16.5 |
| RXRα ( NR2B1) | 28 | 23.4 |
| RXRβ (NR2B2)* | 56 | 5.3 |
| RXRγ (NR2B3) | n/a | 0 |
| THRA (NR1A1) | 41 | 18.2 |
| THRB (NR1A2) | 2 | 3 |
| LXRA (NR1H3) | 9 | 6.4 |
| LXRB (NR1H2) | 50 | 44.8 |
| COUP-TF1 (NR2F2) | 3 | 2.4 |
| COUP-TF2 (NR2F2) | 34 | 39.7 |
| VDR (NR1I1) | 9 | 7.3 |
| FXR (NR1H4) | n/a | 0.2 |
| PPARA | 14 | 7.7 |
| PPARD | 35 | 17.2 |
| PPARG | 7 | 11.4 |
| NURR (NRA42) | 2 | 0.9 |
| PXR (NR1I2) | 0.6 | 0 |
| CAR (NR1I3) | 26 | 0.3 |

*TPM = Transcripts per kilobase million, TPM values reported by the European Bioinformatics Institute Expression Atlas (ebi.ac.uk) and Human Protein Data Bank (proteinatlas.org) for Type II nuclear receptors (T2NRs) expression in U2OS cells. *Note: We also observe expression of RXRβ in U2OS cells when probed with RXR antibody specific to the C-terminal epitope (Proteintech, #212181AP) in western blot (Figure S2B)*

**Table S2.** Plasmid constructs used for co-immunoprecipitation and to make stable cell lines.

| Name | Promoter | Gene product | Short name in the paper | Appeared in |
|---|---|---|---|---|
| PB EF1α RARα-GDGAGLIN-Halo-3xFLAG IRES Puro | EF1α | RARα C-terminally fused with 3xFLAG-Halo tag through a short peptide linker sequence (GDGAGLIN) | RARα-Halo | Figure 2 (A-C), Figure S4 (A-C), Figure S6A |
| PB EF1α V5-Halo-GDGAGLIN-RXRα IRES Puro | EF1α | RXRα N-terminally fused with V5-Halo tag through a short peptide linker sequence (GDGAGLIN) | Halo-RXRα | Figure 2 (A-C), Figure S2C, Figure S4(A-C), Figure S6A |
| PB EF1α RARα_M379R_T386R-GDGAGLIN-Halo-3xF IRES Puro | EF1α | $RAR^{RR}α$ C-terminally fused with 3xFLAG-Halo tag through a short peptide linker sequence (GDGAGLIN) | $RARα^{RR}$-Halo | Figure S5A |
| PB Puro EF1α-RARα-GDGAGLIN-3xFLAG | EF1a | RARα C-terminally fused with 3xFLAG tag through a short peptide linker sequence (GDGAGLIN) | RARα | Figure S2C |
| PB EF1a V5-RXRα IRES Puro | EF1α | RXRα N-terminally fused with V5 tag through a short peptide linker sequence (GDGAGLIN) | RXRα | Figure S2C, Figure S5A |
| PB EF1α ExMCS-GDGAGLIN-SNAPf-3xFLAG IRES Puro | EF1α | SNAPf-3xFLAG tag with a short peptide linker sequence (GDGAGLIN) | SNAP | Figure 2 (D-F), Figure S5(B-D) Figure S6 (B & C), Figure S8 (A & B) |
| PB EF1α RARα-GDGAGLIN-SNAPf-3xFLAG IRES Puro | EF1α | RARα C-terminally fused with 3xFLAG-SNAPf tag through a short peptide linker sequence (GDGAGLIN) | RARα-SNAP | Figure 2 (D-F), Figure 3 (B &C), Figure S5(B-D), Figure S6 (B & C), Figure S8 (A & B) |
| PB EF1α $RAR^{RR}α$ -GDGAGLIN-SNAPf-3xFLAG IRES Puro | EF1α | $RAR^{RR}α$ C-terminally fused with 3xFLAG-SNAPf tag through a short peptide linker sequence (GDGAGLIN) | $RARα^{RR}$-SNAP | Figure 2 (D-F), Figure 3 (B &C), Figure S5(B-D), Figure S6 (B & C), Figure S8 (A & B) |
| PB EF1α V5-SNAPf-GDGAGLIN-RXRα IRES Neo | EF1α | RXRα N-terminally fused with V5-SNAPf tag through a short peptide linker sequence (GDGAGLIN) | SNAP-RXRα | Figure 2 (D-F), Figure 3 (B &C), Figure S5(B-D), Figure S6 (B & C), Figure S8 (A & B) |
| PB PGK Puro EF1α SNAPf- 3xNLS | EF1α | SNAPf -3xNLS linked through a short linker peptide sequence (GDGAGLIN) | SNAP-NLS | Figure 3B, Figure S8 (A & B) |

**Table S3.** Summary of SMT analysis for all conditions.

| Name | $f_{bound}$ | stdev | Num of cells | #Trajs | Appeared in |
|---|---|---|---|---|---|
| H2B-Halo | 75.5 | 0.7 | 22 | 108955 | Figure 1 (C and D) |
| Halo-3xNLS | 11 | 0.9 | 22 | 67555 | Figure 1 (C and D) |
| K. I RARα.c156 | 49 | 1 | 22 | 100148 | Figure 1 (C and D)<br>Figure 2 (C and F)<br>Figure S7 (A and B) |
| K. I RARα.c239 | 47 | 1 | 22 | 42405 | Figure 1 (C and D)<br>Figure 2C |
| K. I RARα.c258 | 53 | 1 | 22 | 74195 | Figure 1 (C and D)<br>Figure 2C |
| K. I RXRα.c10 | 36 | 1 | 20 | 29690 | Figure 1 (C and D)<br>Figure 2C |
| K. I RXRα.D6 | 36 | 1 | 20 | 24886 | Figure 1 (C and D)<br>Figure 2 (C and F)<br>Figure S7 (A and B) |
| K. I RXRα.D9 | 42 | 1 | 20 | 30594 | Figure 1 (C and D)<br>Figure 2C |
| K. I RARα.c156 w/o atRA | 53 | 1 | 22 | 14679 | Figure S3A |
| K. I RARα.c156 w/ 1nM atRA | 54 | 1 | 22 | 15096 | Figure S3A |
| K. I RARα.c156 w/ 100nM atRA | 53 | 1 | 22 | 13795 | Figure S3A |
| K. I RXRα.D6 w/o atRA | 34 | 1 | 20 | 17495 | Figure S3A |
| K. I RARα.D6 w/ 1nM atRA | 31 | 1 | 20 | 17358 | Figure S3A |
| K. I RARα.D6 w/ 100nM atRA | 31 | 1 | 20 | 18899 | Figure S3A |
| O.E RARα-Halo | 27 | 0.72 | 22 | 21997 | Figure 2C<br>Figure S4 (B and C) |
| O.E Halo-RXRα | 11 | 0.75 | 22 | 62767 | Figure 2C<br>Figure S4 (B and C) |
| K. I RARα w/ SNAP | 47 | 1 | 22 | 35396 | Figure 2F<br>Figure S7 (A and B) |
| K. I RARα w/ RARα$^{RR}$ SNAP | 43 | 4 | 43 | 46435 | Figure 2F<br>Figure S7 (A and B) |
| K. I RARα w/ SNAP-RXRα | 49 | 1 | 22 | 41411 | Figure 2F<br>Figure S7 (A and B) |
| K. I RARα w/ RARα-SNAP | 29 | 5 | 46 | 34378 | Figure 2F<br>Figure S7 (A and B) |
| K. I RXRα w/ SNAP | 38 | 1 | 22 | 26031 | Figure 2F<br>Figure S7 (A and B) |
| K. I RXRα w/ RARα$^{RR}$-SNAP | 35 | 1 | 20 | 24137 | Figure 2F<br>Figure S7 (A and B) |
| K. I RXRα w/ SNAP-RXRα | 16 | 2 | 22 | 24910 | Figure 2F<br>Figure S7 (A and B) |
| K. I RXRα w/ RARα-SNAP | 56 | 1 | 22 | 25373 | Figure 2F<br>Figure S7 (A and B) |

*#trajs refers to total number of trajectories of the whole set.*

**Table S4.** Number of molecules estimated according to previously published protocol (Cattoglio et al., 2019).

| **Name** | **Est. # molecules** | **Appeared in** |
|---|---|---|
| Negative control | 0 (fixed) | Figure S4A<br>Figure S5B |
| CTCF | 109,800 (standard) | Figure S4A<br>Figure S5B |
| K. I RARα.c156 | 57024±7048 | Figure 2B (replicate 1) and 2E<br>Figure S4A |
| K. I RARα.c239 | 51596±6144 | Figure 2B (replicate 2) |
| K. I RARα.c258 | 56516±5260 | Figure 2B (replicate 3) |
| K. I RXRα.c10 | 53548±4042 | Figure 2B (replicate 1) |
| K. I RXRα.D6 | 50702±5616 | Figure 2B (replicate 2) and 2E<br>Figure S4A |
| K. I RXRα.D9 | 50946±5645 | Figure 2B (replicate 3) |
| K. I RARα (average) | 55045±2997 | Figure 2B |
| K. I RXRα (average) | 51732±1577 | Figure 2B |
| K. I RARα.c156 w/o atRA | 52586±2230 | Figure S3A |
| K. I RARα.c156 w/ 1nM atRA | 39609±3180 | Figure S3A |
| K. I RARα.c156 w/ 100nM atRA | 33593±4673 | Figure S3A |
| K. I RXRα.D6 w/o atRA | 47130±3291 | Figure S3A |
| K. I RXRα.D6 w/ 1nM atRA | 45849±660 | Figure S3A |
| K. I RXRα.D6 w/ 100nM atRA | 43021±1946 | Figure S3A |
| O.E RARα-Halo | 233732±3738 | Figure 2B<br>Figure S4A |
| O.E Halo-RXRα | 905005±1464 | Figure 2B<br>Figure S4A |
| K. I RARα w/ SNAP | 65617±1293 | Figure 2F<br>Figure S5B |
| K. I RARα w/ RARα$^{RR}$-SNAP | 52532±835 | Figure 2F<br>Figure S5B |
| K. I RARα w/ SNAP-RXRα | 62392±1532 | Figure 2F<br>Figure S5B |
| K. I RARα w/ RARα-SNAP | 37255±371 | Figure 2F<br>Figure S5B |
| K. I RXRα w/ SNAP | 50898±1791 | Figure 2F<br>Figure S5B |
| K. I RXRα w/ RARα$^{RR}$-SNAP | 45148±106 | Figure 2F<br>Figure S5B |
| K. I RXRα w/ SNAP-RXRα | 42248±528 | Figure 2F<br>Figure S5B |
| K. I RXRα w/ RARα-SNAP | 58880±1462 | Figure 2F<br>Figure S5B |

*Errors were calculated from three biological replicates.*
